# Human Retinal Ganglion Cells Respond to Evolutionarily Conserved Chemotropic Cues for Intra Retinal Guidance and Regeneration

**DOI:** 10.1101/2023.02.01.526677

**Authors:** Murali Subramani, Matthew Van Hook, Mohanapriya Rajamoorthy, Fang Qiu, Iqbal Ahmad

**Author notes:** **Correspondence:** Iqbal Ahmad, Ph.D., Department of Ophthalmology and Visual Science, University of Nebraska medical Center, 985840 Nebraska Medical Center, Omaha, Nebraska 68198, USA.

## Abstract

Retinal ganglion cells (RGCs) connect the retina with the higher centers in the brain for visual perception. Their degeneration leads to irreversible vision loss in glaucoma patients. Since human RGCs (hRGCs) are born during fetal development and connections with the central targets are established before birth, the mechanism underlying their axon growth and guidance remains poorly understood. Here, using RGCs directly generated from human embryonic stem cells, we demonstrate that hRGCs express a battery of guidance receptors. These receptors allow hRGCs to read the spatially arrayed chemotropic cues in the developing rat retina for the centripetal orientation of axons toward the optic disc, suggesting that the mechanism of intra-retinal guidance is conserved in hRGCs. The centripetal orientation of hRGCs axons is not only in response to chemo-repulsion but also involves chemo-attraction, mediated by Netrin-1/DCC interactions. The spatially arrayed chemotropic cues differentially influence hRGCs physiological responses, suggesting that neural activity of hRGCs may facilitate axon growth during inter-retinal guidance. Additionally, we demonstrate that Netrin-1/DCC interactions, besides promoting axon growth, facilitate hRGCs axon regeneration by recruiting the mTOR signaling pathway. The diverse influence of Netrin-1/DCC interactions ranging from axon growth to regeneration may involve recruitment of multiple intracellular signaling pathways as revealed by transcriptome analysis of hRGCs. From the perspective of *ex-vivo* stem cell approach to glaucomatous degeneration, our findings posit that *ex-vivo* generated human RGCs are capable of reading the intra-retinal cues for guidance toward the optic disc, the first step toward connecting with the central target to restore vision.

## INTRODUCTION

Retinal ganglion cells (RGCs) are the main output neurons that wire the retina with higher centers of the brain for the perception of vision. Degeneration of these cells in glaucoma patients disrupts this wiring leading to irreversible loss. E*x-vivo* stem cell therapy for glaucomatous degeneration is a practical approach whose success depends upon transplanted human RGCs’ ability to elaborate axons that can read guidance cues and connect properly with the central targets. The guidance of RGC axons to target is a complex process, which consists of several stages that include orientation of axons within the retina for their proper exit, guidance without the retina at the optic chiasm, and finally making retinotopic connections with the cells in the superior colliculus (SC) in the midbrain and lateral geniculate nucleus (LGN) in the thalamus (Harada et al., 2007). Significant progress has been made in understanding the mechanism involving evolutionary conserved molecules and receptors that guide RGC axons at each of the three stages of wiring with the brain (Erskine and Herrera, 2014). However, whether or not the hRGCs recruit similar mechanisms to find the central targets is not well understood. Here, we have examined if hRGCs are capable of reading the cues within the retina to orient their axons toward the optic disc for a proper exit, the first step toward connecting with the central targets.

Evidence suggests that the centripetal orientation of RGC axons toward the optic disc is regulated by the chemo-repulsive environment in the peripheral retina, which constraints them toward the chemo-attractive center, away from the periphery (Snow et al., 1991). The chemo-repulsive environment is likely due to the expression of Slit proteins (Niclou et al., 2000, Thompson et al., 2006, Thompson et al., 2009) and chondroitin sulfate proteoglycans (CSPGs) (Brittis et al., 1992, Brittis and Silver, 1995). Slits (Slit1-3 in mammals) are highly conserved secretory proteins, which mediate chemo-repulsion through transmembrane receptors, ROBO (ROBO1-4 in mammals), an acronym from the *Drosophila* mutant ROundaBOut (Blockus and Chedotal, 2016). *Slit1-2* and *Robo1-2* are expressed in the developing retina, with a central to peripheral gradient of expression for the former (Ringstedt et al., 2000). The boundary of their expression progressively shifts toward the periphery, constraining the growth of the newly differentiating RGC axons toward the center. Loss of Slit proteins or ROBO (Thompson et al., 2006, Thompson et al., 2009) and enzymatic digestion of CSPG disrupt the projection and cause aberrant growth of RGC axons (Brittis et al., 1992). CSPGs are produced by developing and adult neurons and the major CSPG types in the CNS include leticans, phosphocans and NG2 (Galtrey and Fawcett, 2007, Dyck and Karimi-Abdolrezaee, 2015, Bandtlow and Zimmermann, 2000).

The chemo-attractive environment in the central retina for the centripetal orientation of RGC axons is less well defined although evidence suggests that Netrins may be involved. Netrins (Netrin-1 and 2 in mammals) are highly conserved bidirectional guidance proteins, attracting growth cones through the transmembrane receptor deleted in colorector cancer (DCC), or Neogenin and repelling them via a transmembrane receptor, UNC5A-D, an acronym for the *c. elegans* mutant phenotype, UNCoordinated (Lai Wing Sun et al., 2011). Netrin-1 is expressed predominantly in the optic nerve head and Netrin-1/DCC interactions serve as a chemoattractant for RGC neurites (Deiner et al., 1997). Two guidance-related retinal phenotypes were observed in Netrin-1 deficient mice; the RGCs axons failed to enter the optic disc and though centrally polarized, the axons grew directionless into other retinal regions (Deiner et al., 1997). These results demonstrated that Netrin-1 is required for the guidance of RGC axons out of the eyes that requires their proper and directed orientation toward the optic disc.

In order to examine the intra-retinal guidance potential of hRGC axons, we tested a hypothesis that hRGCs can recognize and respond to evolutionarily conserved and spatially distributed chemotropic cues in the developing rat retina. The premise was tested by co-culturing human embryonic stem cells derived RGCs on central and peripheral embryonic day 16 (E16) rat retinal cells. We observed that the dimension of hRGCs neurites was remarkably different on central versus peripheral embryonic rat retinal cells. The complexity of neurites and the length of axons were significantly greater in the former compared to the latter condition, regulated by Netrin-1/DCC and Slits/ROBO2 plus CSPG interactions, respectively, suggesting that hRGCs respond to spatially distributed chemotropic cues in the embryonic retina that facilitate intra-retinal guidance of RGC axons. The spatially arrayed chemotropic cues, in addition to influencing neurite morphology and axon growth, affect the neural activity of hRGCs as revealed by electrophysiological responses, regulated by Netrin-1/DCC and Slits/ROBO2intercations and by CSPG as well. Given the emerging role of neural activity in axon growth (Fields, 1994, Hanson and Landmesser, 2004, McLaughlin et al., 2003, Nicol et al., 2007) it may be surmised that chemotropic cues-mediated neural activity influences hRGC axon growth. As surrogates for central and peripheral RGCs, we examined the axon regeneration potential of hRGCs; we observed that, when co-cultured with E16 central rat retinal cells, hRGCs regenerated their axons more efficiently following axotomy than those co-cultured with E16 peripheral rat retinal cells. The preferential axon regeneration in the former was facilitated by Netrin-1/DCC interactions and was dependent on mTOR signaling. Transcriptome analysis of hRGCs co-cultured with E16 central retinal rat cells where Netrin-1/DCC interactions were blocked through neutralization by anti-DCC antibody revealed the recruitment of multiple intercellular pathways, which are known to influence diverse Netrin-mediated functions including axon growth, neural activity, and regeneration. Together, our results demonstrate for the first time the behavior of nascent hRGCs vis-à-vis chemotropic cues in the backdrop of information available from animal models, predicting that the early events guiding hRGCs’ axons to the optic disc in the developing human retina are evolutionarily conserved. From the perspective of *ex-vivo* stem cell approach to glaucomatous degeneration, our findings posit that *ex-vivo* generated human RGCs are capable of reading the intra-retinal cues for guidance toward the optic disc, the first step toward connecting with the central target to restore vision.

## RESULTS

### Human RGCs express functional guidance receptors

To examine the axon guidance potential of hRGCs, we directly generated RGCs from a genome engineered human BRN3b-reporter ESC line (Sluch et al., 2017), hESC^Brn3b-tDT^ our stage-specific chemically defined protocol that recapitulates the developmental mechanism (**Fig. 1A**). The method was modified for generating retinal progenitor cells (RPCs) using high-density monolayer culture of ES cells for batch-to-batch consistency, which may be compromised in embryoid bodies-based method (Teotia et al., 2017a). The differentiation of hESC^Brn3b-tDT^ along RGC lineage activated *BRN3B* promoter leading to the expression of tdTomato (tdT) fluorescent protein as an RGC reporter (Sluch et al., 2017). The proportion of directly differentiated nascent (tdT^+^ β-tubulin^+^ cells) and matured (tdT^+^ PAX6^+^ cells) was 53.68±4%, and 41.89±4%, respectively (**Fig. 1B,C**). That the differentiation of RGCs was regulated and not random was demonstrated by the temporal induction of the genes corresponding to RGC regulators (*ATOH7, BRN3B* and *ISLET1)* and markers (*SNCG* and *THY1*) (**Fig. 1D**). The guidance of RGC axons within, and without the retina toward the target is facilitated by a battery of chemotropic guidance molecules/cues (=chemotropic cues) recognized by receptors on the RGC growth cones (Harada et al., 2007). Therefore, we examined whether or not the *ex-vivo* generated hRGCs have the capability to recognize the guidance cues by their expression of the cognate receptors. We observed temporal activation of transcripts corresponding to *DCC, ROBO2, NRP1*, and *EPHs*, receptors that recognize chemotropic molecules *NETRIN-1, SLIT2, SEMAPHORIN*, and different *EPHRIN*, respectively during hRGC differentiation (**Fig. 2A**). The temporal increase in their expression suggested a regulated activation of these genes during RGC generation. Next, to determine the functional status of the chemotropic receptors, hRGCs^Brn3b-tDT^ were cultured in the soma chamber of a microfluidic device and activation/collapse of growth cones (GCs), as determined by the number of filopodia, was examined in the presence of chemotropic cues in the axon chamber. We observed that compared to controls, GCs were activated in the presence of Netrin-1, whose effect was abrogated in the presence anti-DCC antibody, suggesting a functional Netrin-1/DCC chemo-attractive interactions (**Fig. 2B**). In contrast, exposure to chemo repellants, Slit2 (**Fig. 2C**) or chondroitin sulfate proteoglycans (CSPG) (**Fig. 2D**) caused GC collapse, compared to controls. Antibody-mediated neutralization of ROBO2 or degradation of CSPG by chondroitinase restored the activated status of GCs. Together, these observations demonstrated that *ex-vivo* generated hRGCs express functional axon guidance receptors, suggesting their capacity to respond intra-retinal guidance cues in the developing rat retina.

**Fig. 1.**
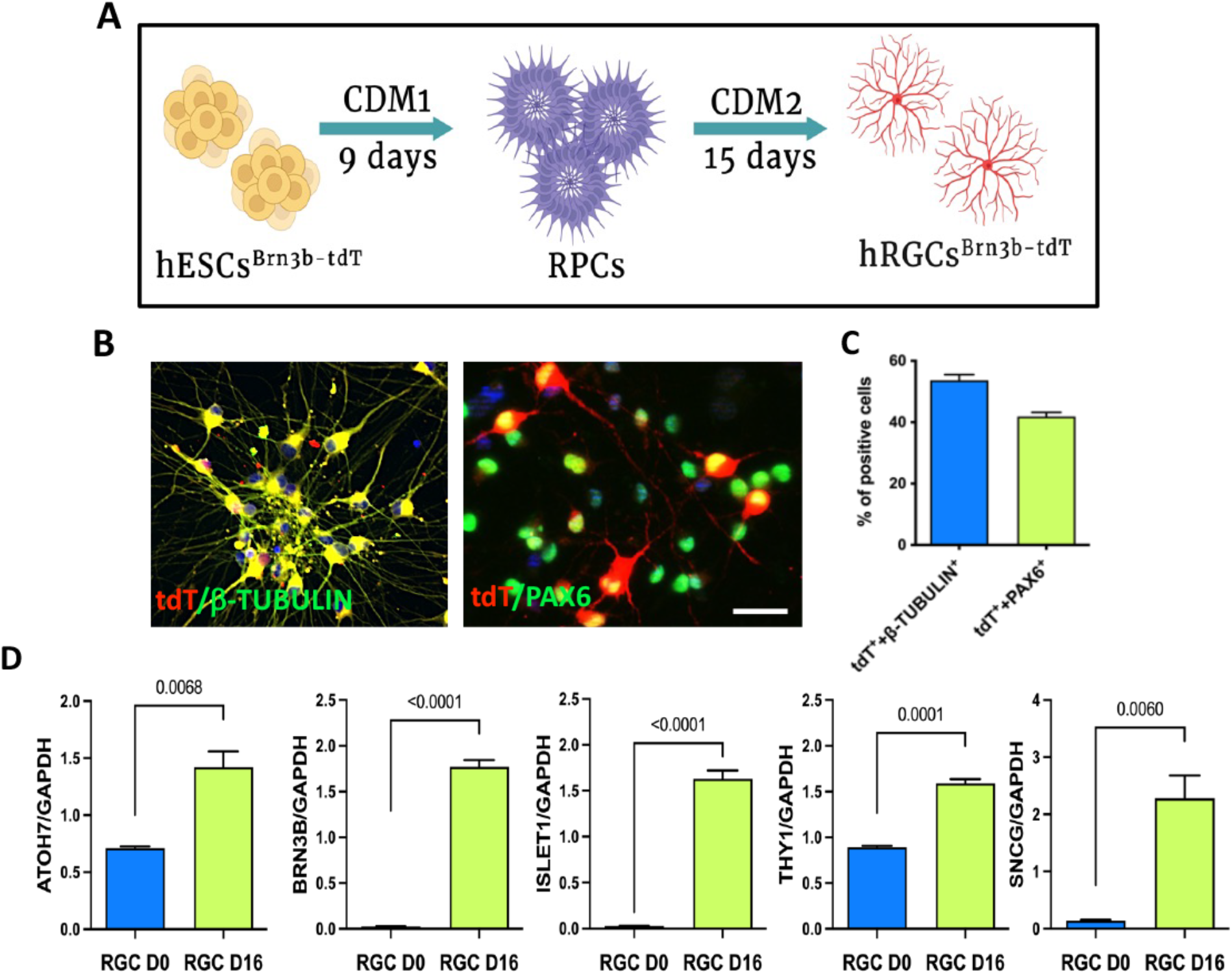
Direct generation of hRGCs from human BRN3b-reporter ESC line by recapitulating developmental mechanism: **A**. A schematic representation of 2D method of hRGC generation from hESC through the retinal progenitor cells stage. **B**. Representative merged immunofluorescence images showing tdT^+^ hRGCs co-expressing nascent and mature RGC markers β-tubulin and PAX6 respectively. **C**. Proportion of hRGCs expressing nascent (tdT^+^ β-tubulin^+^) and mature (tdT^+^ Pax6^+^) markers. **D**. A temporal qPCR analysis showing regulated activation of genes corresponding to hRGC regulators (*ATOH7, BRN3B, ISLET1*) and markers (*THY1, SNCG*) at DIV 16 vs DIV 0. Values are expressed as mean±s.e.m. (Student’s t-test; P<0.05). Experiments were carried out in triplicates per group. Scale bars: 20 μm. CDM=Chemically defined medium (see methods). DIV=Day *in vitro*.

**Fig. 2.**
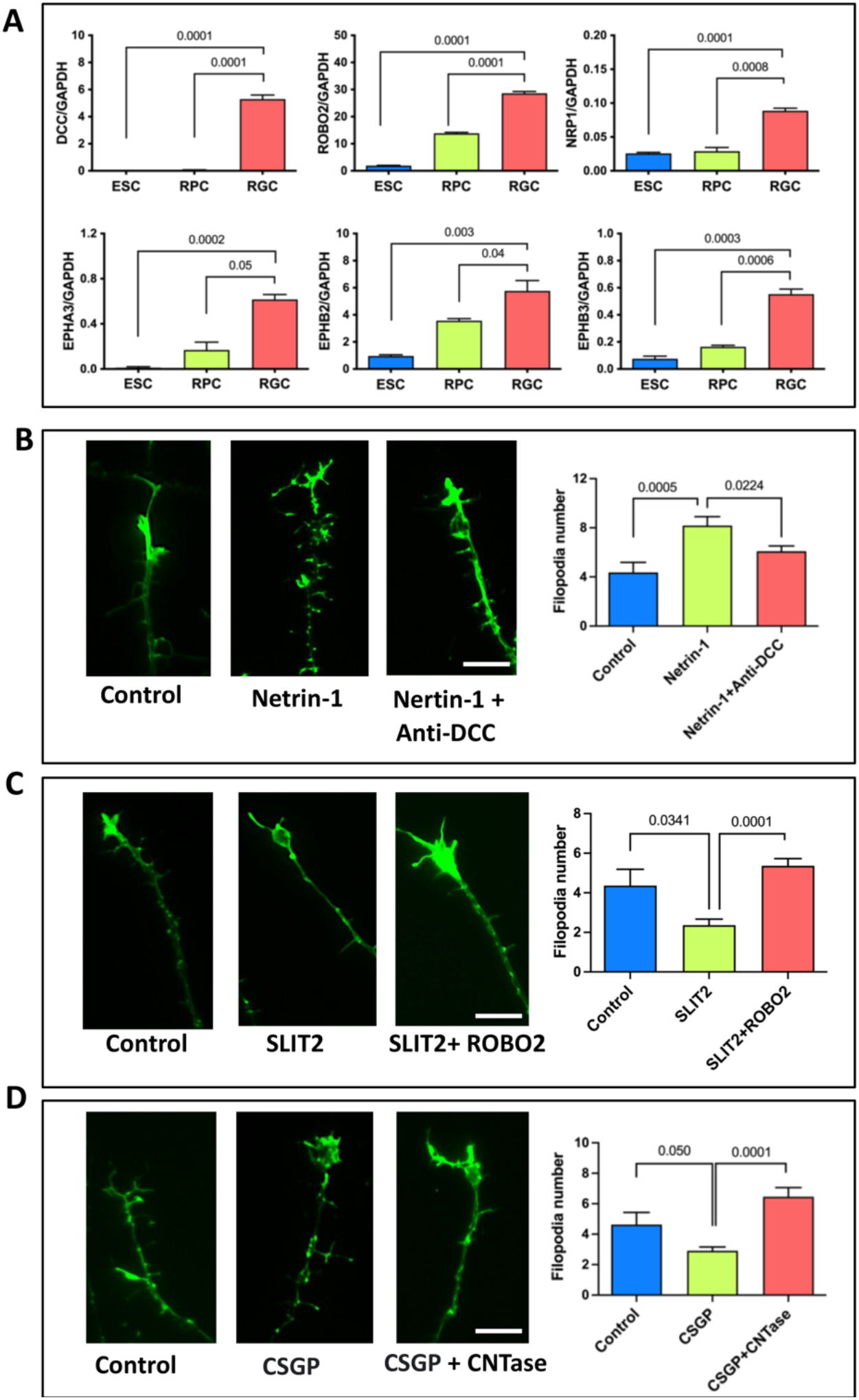
Expression of functional guidance receptors in hRGCs: **A**. A temporal qPCR analysis reveals a regulated activation of genes corresponding to guidance receptors, *DCC, ROBO2, NRP1, EphA3, EphB2*, and *EphB3* during hRGC differentiation. **B-D**. Phalloidin-immunofluorescence images of hRGCs growth cones showing filopodial number in the presence Netrin-1 and Netrin-1 plus DCC antibody (B); Slit2 and Slit plus ROBO2 antibody (C), and CSPG and CSPG plus chondroitinase (CNTase) (C). Values are expressed as mean±s.e.m. (Student’s t-test; P<0.05, ns, not significant). Experiments were carried out in triplicates per group. Scale bars: 20 μm.

### Human RGCs axons respond to spatially distributed chemotropic cues in the developing rat retina

Evidence suggests that chemotropic cues are differentially distributed across the central versus peripheral developing rodent retina for centripetal orientation of RGC axons toward the optic disc (Brittis et al., 1992). To examined whether hRGCs could respond to these spatial chemotropic cues hRGCs^Brn3b-tDT^ were co-cultured on cells dissociated from embryonic day 16 (E16) central and peripheral retina (**Fig. 3A**). E16 represents the peak of RGC generation in rats, when the wave of RGC differentiation spread out from the central to peripheral retina (Rapaport et al., 2004). We observed that hRGCs^Brn3b-tDT^ cultured on the central versus peripheral rat retinal cells displayed different morphology; while those on the central retinal cells had long and complex neurites, those on the peripheral retinal cells, in contrast, elaborated shorter and simpler neurites (**Fig. 3B-E**). Examination of hRGCs^Brn3b-tDT^ revealed more collapsed GCs in those co-cultured with the peripheral rat retinal cells than on the central rat retinal cells (**Fig. S1A-C**). The differences in complexity and length of hRGCs ^Brn3b-tDT^ neurites suggested that the central retinal cells offer a more conducive extracellular environment for neuritogenesis and growth of axons than the peripheral retinal cells, which is likely due to spatially distributed chemo-attractant (Netrin-1) and chemo-repulsive cues (Slits and CSPG), respectively. To test this premise, we disrupted the chemotropic signals between the rat cells and hRGCs by antibody-mediated neutralization of specific-chemotropic receptors on hRGCs^Brn3b-tDT^ before co-culture on the central versus peripheral rat retinal cells. The peripheral rat retinal cells were pre-treated with chondroitinase to degrade CSPG before co-culturing with hRGCs^Brn3b-tDT^ in order to determine the influence of CSPG on hRGCs. The tdT-positive neurites were co-stained for SMI32+Tau1 immunoreactivities to examine the effects specifically on the hRGC axons. We observed that both the complexity and lengths of hRGC axons co-cultured with the central rat retinal cells decreased significantly in the group pre-treated with the anti-DCC antibody, compared to the IgG-treated hRGCs^Brn3b-tDT^ controls (**Fig. 4A-D**), demonstrating a facilitatory role of Netrin-1/DCC interactions on hRGC^Brn3b-tDT^ axon growth. In contrast, the complexity and length of hRGC axons co-cultured with the peripheral rat retinal cells increased significantly in the group pre-treated with the anti-ROBO2 antibody, compared to the IgG-treated hRGCs^Brn3b-tDT^ controls (**Figure 4E-H**). Similar results were obtained when CSPG in the peripheral rat retinal cells were degraded by chondroitinase, demonstrating that the inhibitory influence mediated by Slit/ROBO2 interactions or CSPG keep the hRGC ^Brn3b-tDT^ axon growth inhibited (**Figure 4I-L**). The blocking of Netrin-1/DCC interactions in hRGC^Brn3b-tDT^ co-cultured with the central rat retinal cells (**Fig. S2**) and Slits/ROBO2, and CSPG interactions in those co-cultured with the peripheral rat retinal cells (**Fig. S3**) had similar inhibitory and facilitatory effects, respectively, on overall neurite complexity and lengths as those observed for the axons. Together, these observations suggested that hRGCs can detect and respond accordingly to spatially distributed chemo-attractive and chemo-repulsive cues in the developing rat retina.

**Fig. 3.**
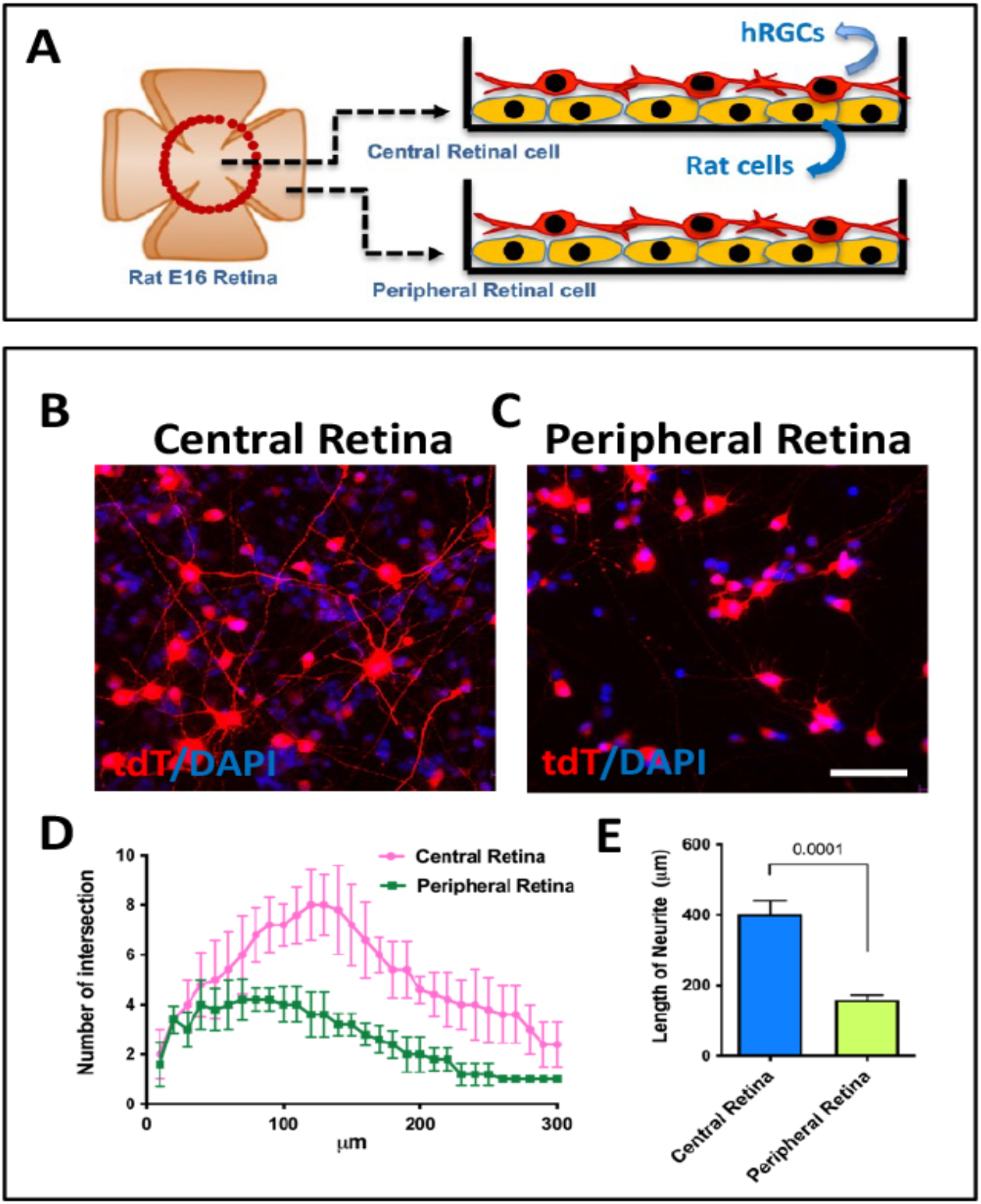
Response of hRGCs axons to spatially arrayed chemotropic cues in the developing central and peripheral rat retina: **A**. A schematic representation of hRGCs^Brn3b-tDT^ and developing central versus peripheral E16 rat retinal cells co-culture paradigm to examine the response to chemotropic cues. **B-E**. hRGCs^Brn3b-tDT^ co-cultured on the central rat retinal cells display complex and longer neurites (B, D, E) than those co-cultured on the peripheral rat retinal cells (C, D, E). Neurite complexity and length was analyzed by Sholl analysis (D, E). Values are expressed as mean±s.e.m. (Student’s t-test; P<0.05, ns, not significant). Experiments were carried out in triplicates per group. Scale bars: 50 μm.

**Fig. 4.**
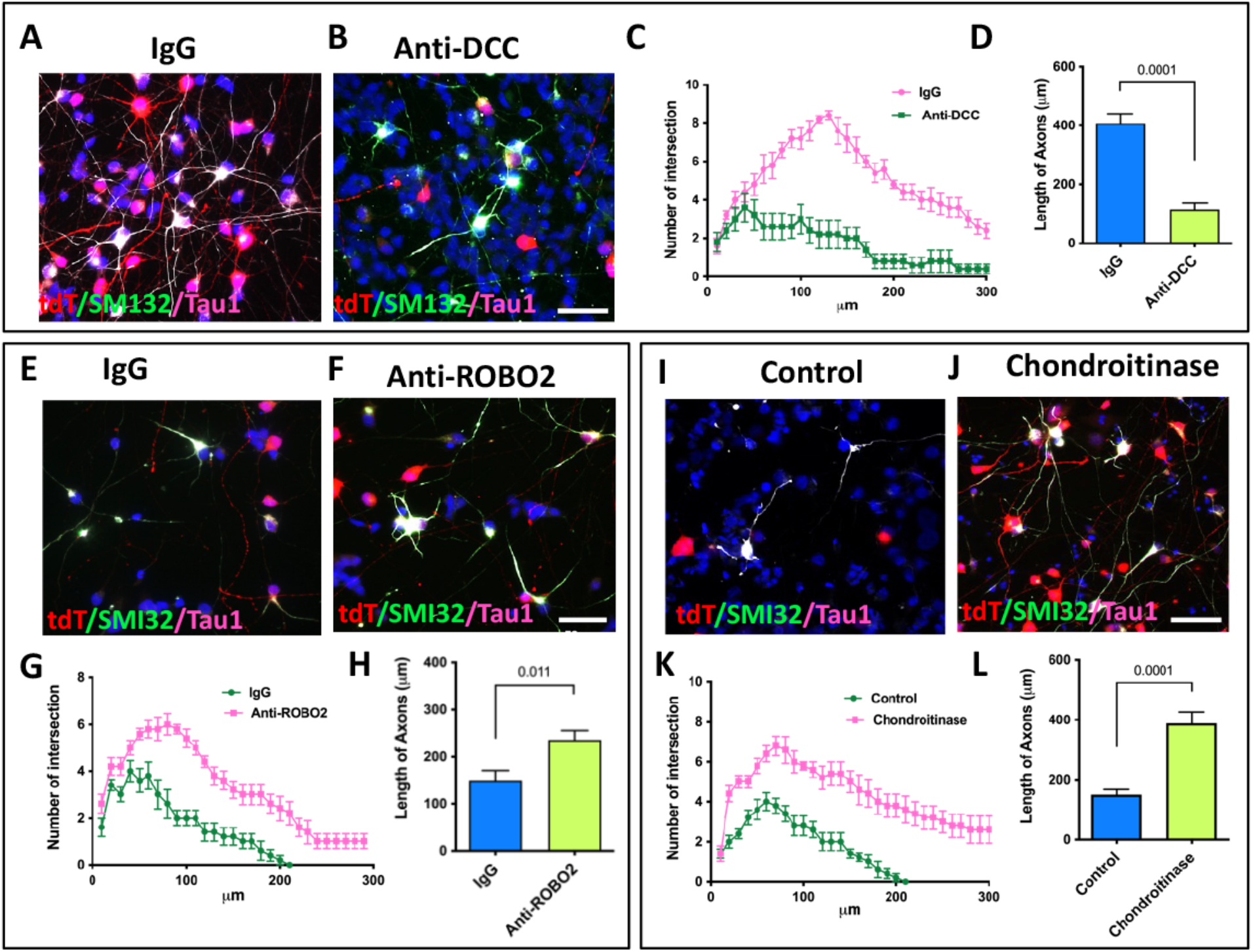
Netrin-1/DCC interactions in mediating chemotropic influence of central rat retinal cells on hRGCs axons: **A-D**. hRGCs^Brn3b-tDT^ preincubated with IgG (A) and DCC-antibody (B) were co-cultured on E16 central retinal cells followed by sholl analyses (C, D) of hRGCs^Brn3b-tDT^ co-expressing SMI32 and Tau immunoreactivities, identifying axons. The complexity of axons and their length were significantly decreased when DCC on hRGCs^Brn3b-tDT^ was neutralized by DCC antibody, compared to IgG controls, suggesting Netrin-1/DCC interactions underlying central retinal cells-mediated chemoattraction. **E-H**. hRGCs^Brn3b-tDT^ preincubated with IgG (E) and ROBO2-antibody (F) were co-cultured on E16 peripheral retinal cells followed by sholl analyses (G,H) of hRGCs^Brn3b-tDT^ co-expressing SMI32 and Tau immunoreactivities, identifying axons. The complexity of axons and their length were significantly increased when Slits on hRGCs^Brn3b-tDT^ was neutralized by ROBO2 antibody, compared to IgG controls, suggesting Slit/ROBO2 interactions underlying peripheral retinal cells-mediated chemo-repulsion. **I-L**. hRGCs^Brn3b-tDT^ were co-cultured on E16 peripheral retinal cells, untreated control (I) and treated with chondroitinase (J) followed by sholl analyses (K, L) of hRGCs^Brn3b-tDT^ axons. The complexity of axons and their length were significantly increased when hRGCs^Brn3b-tDT^ were cultured on cells treated with chondroitinase, compared to untreated controls, suggesting that CSPG distributed on peripheral retinal cells underlie peripheral retinal cells-mediated chemo-repulsion. Values are expressed as mean±s.e.m. (Student’s t-test; P<0.05, ns, not significant). Experiments were carried out in triplicates per group. Scale bars: 50 μm.

### Human RGC axons orient centripetally in response to netrin in the developing rat retina

While the co-culture of hRGCs^Brn3b-tDT^ on the central versus peripheral retinal cells demonstrated the ability of these to cells recognize the spatial chemotropic cues, the approach could not show their influence on the orientation of neurites because the retinal axes were lost due to the dissociation of cells. Therefore, we transplanted hRGCs^Brn3b-tDT^ on PN1 rat central retinal explants away from the optic disc to examine the orientation of the neurites. Examination of the explants five days later by two-photon microscopy revealed the integration of hRGCs^Brn3b-tDT^ in the host retina and orientation of their neurites centrally toward the optic disc (**Fig. 5A**). Both the length and the centripetal orientation of the neurites were compromised when hRGCs^Brn3b-tDT^ were pre-incubated with anti-DCC antibody (**Fig. 5B**). Next, we transplanted hRGCs^Brn3b-tDT^ near the optic disc in PN1 rat retina explants. Analysis of transplants 5 days later revealed both the hRGCs^Brn3b-tDT^ and the neurites preferentially oriented and with the optic disc (**Fig. 5C**), which was abrogated in the presence of anti-DCC antibody (**Fig. 5D**). These results suggested that that the centripetal orientation of hRGC axons is likely due to Netrin-1/DCC chemo-attraction. However, a previous study has demonstrated that Netrin-1 expression in the developing rodent retina is confined to optic disc, whereas our results demonstrated the activity of Netrin-1 in the central rat retinal cells. To address the conflicting observations, we carried out immunocytochemical and qPCR analyses of Netrin-1 expression in the central and peripheral E16 rat retina. Immunocytochemical analysis of E16 rat retina revealed robust Netrin-1 immunoreactivities in the optic disc as previously described. However, a relatively lower level of immunoreactivity was present in the central retina, which tended to decrease centrifugally (**Fig. S4A**). This result was corroborated by qPCR analysis of *Netrin-1* transcripts, whose expression was significantly higher in the central rat retinal cells versus peripheral rat retinal cells (**Fig. S4B**). The presence of Netrin-1 immunoreactivity and transcripts in the central retina, suggested that the centripetal orientation of the developing RGC axons may not be a passive process following chemo-repulsion, but actively facilitated by low levels of Netrin-1 expression in the central retina. Next, to know the temporal maintenance of the chemotropic cues in the embryonic and postnatal retina, we co-cultured hRGCs^Brn3b-tDT^ on the central versus peripheral rat retinal cells obtained from PN1 and P15. We observed similar spatially distributed chemotropic cues in PN1/PN15 retina that could be interpreted by hRGCs^Brn3b-tDT^ in P15 retina as in E16 retina (**Fig. 6A-D**). Together, these results demonstrated that hRGCs^Brn3b-tDT^ can read the evolutionarily conserved intra-retinal chemotropic cues for centripetal orientation toward optic disc and that these cues are functional in postnatal retina. Additionally, our results demonstrated that Netrin-1 in the central retinal cells may play an active role in the centripetal orientation of RGC axons in general.

**Fig. 5.**
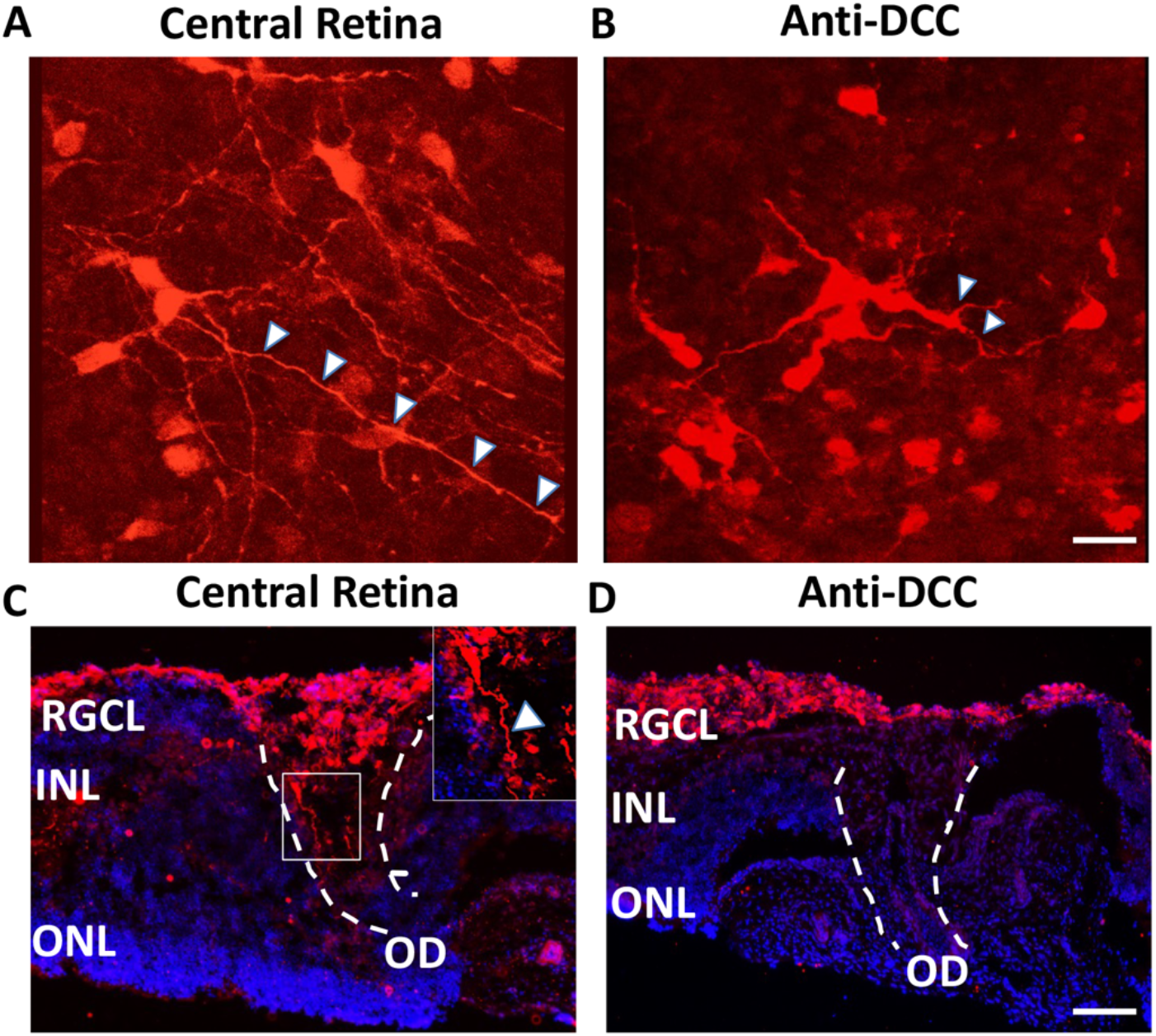
Centripetal orientation of hRGC axons under the influence of Netrin-1/DCC interactions: **A, B**. hRGCs^Brn3b-tDT^, pre-incubated in IgG (A) or DCC antibody (B) were transplanted on PN1 rat retinal explants, away from or on optic disc and examined 5 days later. The centripetal orientation of hRGCs^Brn3b-tDT^ neurites (arrowheads) toward the optic disc was abrogated in the presence of DCC antibody, compared to IgG controls. **C, D**. hRGCs^Brn3b-tDT^, pre-incubated in IgG (C) or DCC antibody (D) were transplanted intravitreally in PN1 rat retina. The penetration of hRGCs^Brn3b-tDT^ processes (arrowhead) into the optic disc was abrogated in the presence of DCC antibody (F), compared to IgG controls (E). n=3 (A-B); n=3 (C-D). Scale bars: 50 μm.

**Fig. 6.**
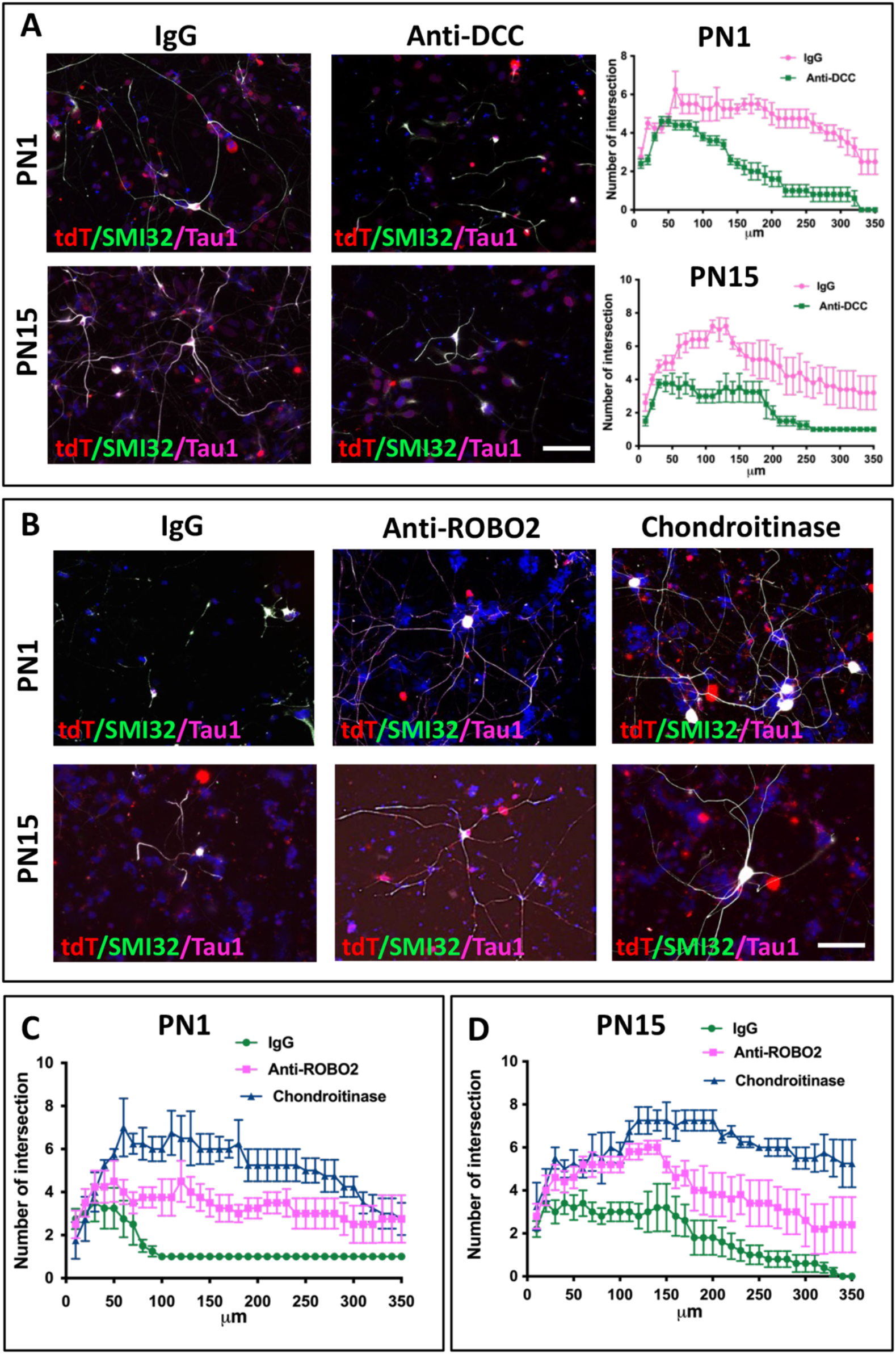
Persistent impact of spatially arrayed chemotropic cues in postnatal rat retina on hRGCs: hRGCs^Brn3b-tDT^ pre-incubated with IgG/anti-DCC antibody/anti-ROBO2 antibody were co-cultured with central and peripheral PN1 (A) and PN15 (B) rat retinal cells untreated or treated with chondroitinase. **A**. Central cells from either PN1 or PN15 retina facilitated the elaboration of hRGCs^Brn3b-tDT^ neurites, which was abrogated when Netrin-1/DCC interaction was blocked. **B**. The elaboration of hRGCs^Brn3b-tDT^ neurites was inhibited on peripheral cells from either PN1 or PN15 retina, which disinhibited when Slits/ROBO2 interactions were blocked or CSPG was degraded. Experiments were carried out in triplicates per group. Scale bars: 50 μm.

### Human RGC physiology is impacted by spatially distributed chemotropic cues

Next, we examined whether or not the spatially distributed chemotropic cues that impact neuritogenesis and axon growth affect the physiology of hRGCs as well. This question is relevant for the following reasons: First, growing evidence suggests that neural activity facilitates axon growth (McLaughlin et al., 2003, Hanson and Landmesser, 2004, Nicol et al., 2007, Fields, 1994). Second, membrane depolarization is known to sensitize the Netrin-1/DCC interactions mediated axon growth in embryonic cortical neurons (Bouchard et al., 2008). Lastly, in adult hippocampal neurons Neterin-1/DCC interactions, besides impacting dendrite and synaptic maturity, has been observed to facilitate long term potentiation (LTP) (Horn et al., 2013). To test this premise, hRGC^Brn3b-tdT^ were cultured on the E16 central (=central hRGC^Brn3b-tdT^; **Fig. 7**) versus peripheral (=Peripheral hRGC^Brn3b-tdT^; **Fig. 8**) rat retinal cells and were monitored by whole cell voltage- and current-clamp recordings. To know if the parameters of voltage-gated Na^+^ currents and spiking activity are influenced by the spatially arrayed chemotropic cues, recordings were compared following the neuralization of their cognate receptors on hRGCs^Brn3b-tdT^ or degradation of CSPG by pre-treating peripheral rat retinal cells with chondroitinase. For example, when recordings were done on the central hRGC^Brn3b-tdT^, pre-treated with DCC neutralizing antibody, the voltage-gated Na^+^ current density was significantly lower than untreated controls, suggesting a role of Netrin-1/DCC interactions in regulating the function of central hRGC^Brn3b-tdT^ (**Fig. 7**). Although the spiking activity in DCC antibody-treated RGCs^Brn3b-tdT^ trended lower than controls, there was no statistically significant difference at any current stimulus, except at 5pA (**Fig. 7D-E**). In contrast, the voltage-gated Na^+^ current density was significantly higher in peripheral hRGC^Brn3b-tdT^, pre-treated with ROBO2 neutralizing antibody or cultured on peripheral rat retinal cells pretreated with chondroitinase, compared to IgG-treated controls, suggesting a role for chemo-repulsive influence on the functional maturity of peripheral hRGC^Brn3b-tdT^. The spiking activity in current-clamp recordings appeared slightly elevated in ROBO2 antibody- and chondroitinase-treated groups, with peak spike number at 12.5 pA stimulus being significantly different between the groups. Together, these observations suggest that hRGCs not only read the spatially distributed chemotropic cues for neuritogenesis and axon polarization, but also respond to them functionally, where centrally distributed Netrin-1 and peripherally distributed Slits and CSPG favor mature and immature functional phenotype, respectively, that may influence axon growth during development and regeneration.

**Fig. 7.**
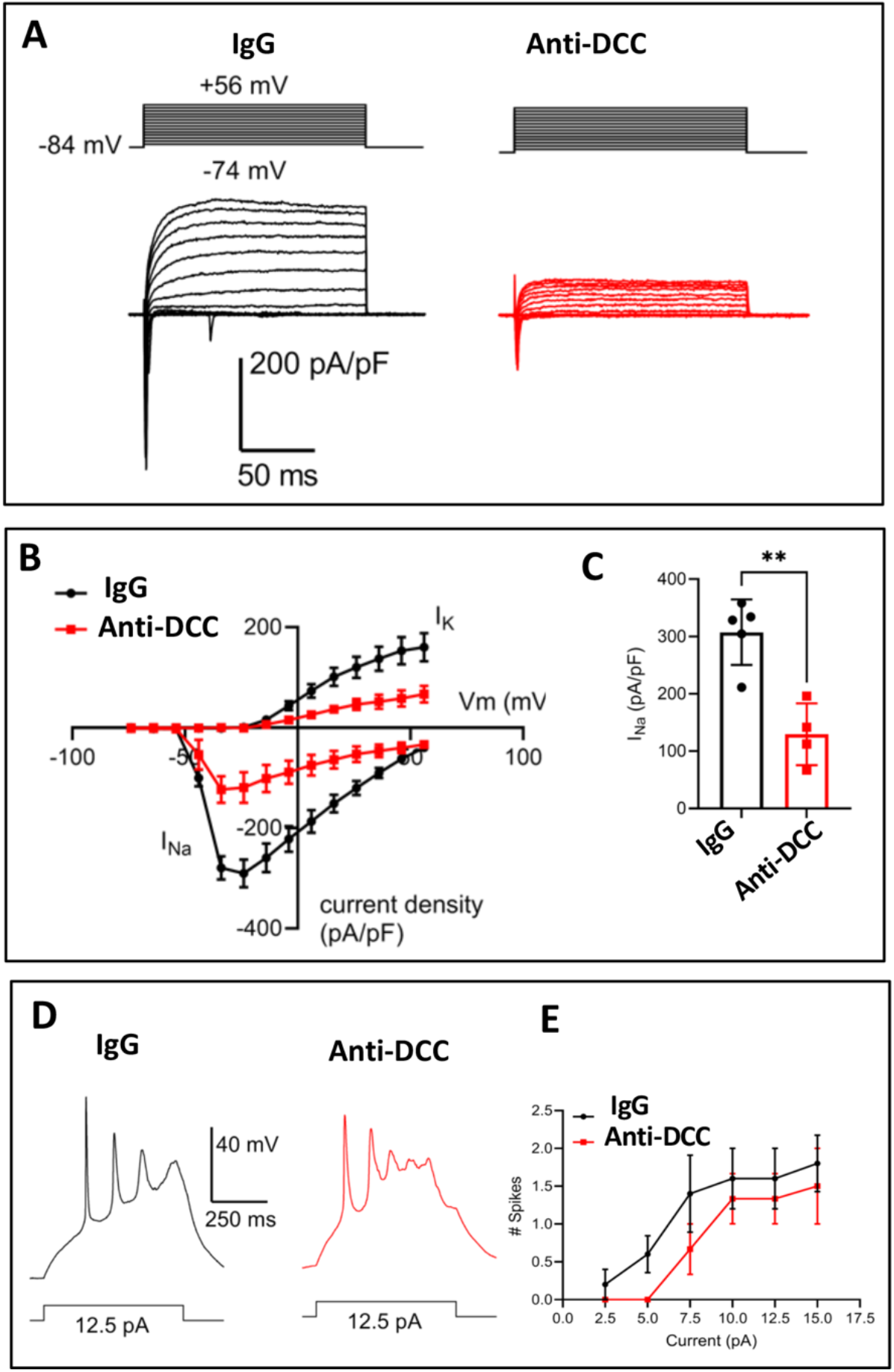
The impact of Nertin-1/DC interactions on hRGC physiology: hRGCs^Brn3b-tDT^, pre-incubated in IgG or DCC antibody were co-cultured on central rat retinal cells following by electrophysiological analysis. **A**. Whole-cell voltage-clamp recording of I_Na_ and I_K_ in response to a series of depolarizing steps from a holding potential of -84 mV (−74 mV to +56 mV, 10 mV increments). **B**. Current-voltage plot of Ina and IK current density. IgG Control n = 5 cells; anti-DCC n = 4 cells. **C**. Peak Na current density. **p=0.002, unpaired t-test. **D**. Example current-clamp traces of spiking in response to a 12.5 pA depolarizing stimulus (500 ms). **E**. Plot of spiking relative to current stimulus.

**Fig. 8.**
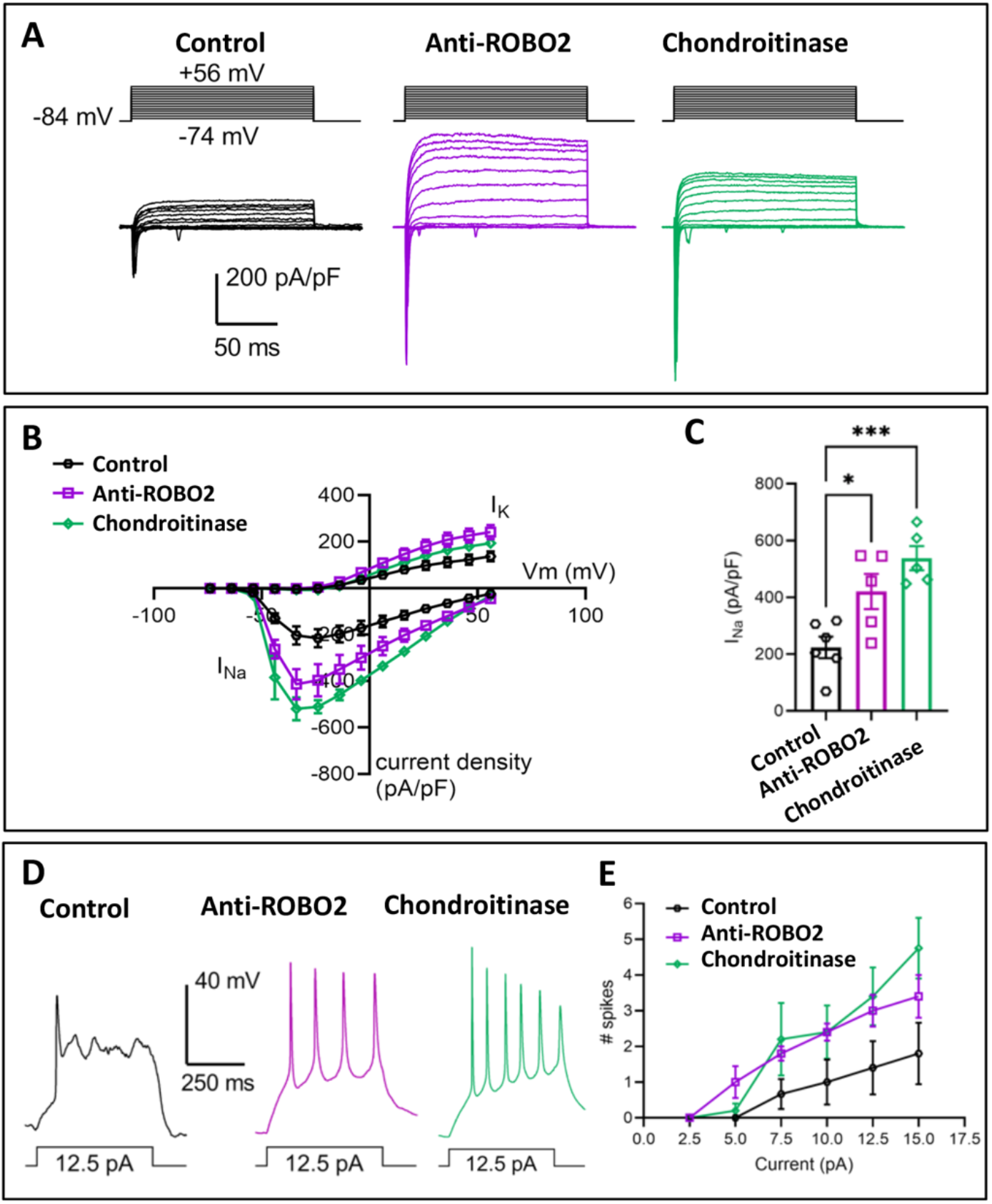
The impact of Slit/ROBO2 interactions and CSPG on hRGC physiology: hRGCs^Brn3b-tDT^, pre-incubated in IgG or ROBO2 antibody were co-cultured on peripheral rat retinal cells following by electrophysiological analysis. Another batch of hRGCs^Brn3b-tDT^ were co-cultured on untreated and chondroitinase-treated peripheral rat retinal cells for electrophysiological analysis. **A**. Whole-cell voltage-clamp recording of I_Na_ and I_K_ in response to a series of depolarizing steps from a holding potential of -84 mV (−74 mV to +56 mV, 10 mV increments). **B**. Current-voltage plot of Ina and IK current density. IgG control n = 6 cells; anti-ROBO2, n = 5 cells; chondroitinase, n = 5 cells. **C**. Peak Na current density. There was a significant difference among groups (p=0.0013) with both anti-ROBO2 and chondroitinase significantly differing from controls *p=0.02; ***p=0.008. **D**. Example current-clamp traces of spiking in response to a 12.5 pA depolarizing stimulus (500 ms). **E**. Plot of spiking relative to current stimulus.

### Human RGC axon regeneration is influenced by chemotropic cues

Next, we examined the influence of the spatially distributed chemotropic cues on the regenerative potential of RGCs, given their influence on axon growth and neural activities of cells, both known to influence the response of cells to injury (Lim et al., 2016). Additionally, in goldfish, where optic nerve regenerates, Netrin 1 and DCC are upregulated after optic nerve injury, suggesting a role for Netrin-1/DCC interactions in regeneration (Petrausch et al., 2000). Therefore, we tested whether axon regeneration following axotomy is enhanced in hRGC^Brn3b-tdT^ cultured in Netrin-1 enriched environment. We cultured hRGC^Brn3b-tdT^ in the soma chambers of the microfluidic devices till their axons grew into all the microgrooves and reached the axon chambers (**Fig. 9A**). The soma chambers were re-seeded with either E16 central or peripheral rat retinal cells, followed by detergent based axotomy of hRGC^Brn3b-tdT^ axons in the axon chamber (Teotia et al., 2019). Five days later, hRGC^Brn3b-tdT^ axons regenerated in axonal chambers in all three groups (**Fig. 9B, C**). However, the number of hRGC axons in the axon chambers were significantly higher in the central retinal cell group versus those in peripheral retinal cell group or the group in which only hRGC^Brn3b-tdT^ were seeded alone as controls (**Fig. 9E**). Although regenerated axon length appeared similar in control and central hRGC^Brn3b-tdT^ groups, it was significantly decreased in the peripheral hRGC^Brn3b-tdT^ group (**Fig.9D**). Together, these observations suggested that the environment of central retinal cells facilitated the regeneration of hRGC^Brn3b-tdT^ axons. Next, to determine the influence of Netrin-1/DCC interactions on the regenerative potential, DCC expressed in hRGC^Brn3b-tdT^ was neutralized using anti-DCC antibody before co-culture with E16 central rat retinal cells (**Fig. 10**). We observed that both the number and length of the regenerated axons were significantly reduced in the anti-DCC group compared the IgG controls (**Fig. 10A, B, E, F**). Next, we examined if Netrin-1/DCC interactions recruited mTOR signaling, the key intracellular pathway involved in the regeneration of RGC axons (Park et al., 2008, Teotia et al., 2019). We observed that the number and length of axons decreased when regeneration were carried out in the presence of rapamycin (**Fig. 10D-G**). The decrease in regeneration was accompanied by a decrease in the number of hRGC^Brn3b-tdT^ cells expressing pS6, a readout of mTOR signaling. That Netrin1/DCC interactions recruit mTOR pathway was further demonstrated by increase and decrease in the levels of pmTOR/pS6 in hRGC^Brn3b-tdT^ cultured in the presence of Netrin-1 alone and after pretreatment with anti-DCC antibody, respectively (**Fig. S5**). Together, these results suggest that Netrin-1/DCC interactions facilitate regeneration in hRGCs via mTOR signaling.

**Fig. 9.**
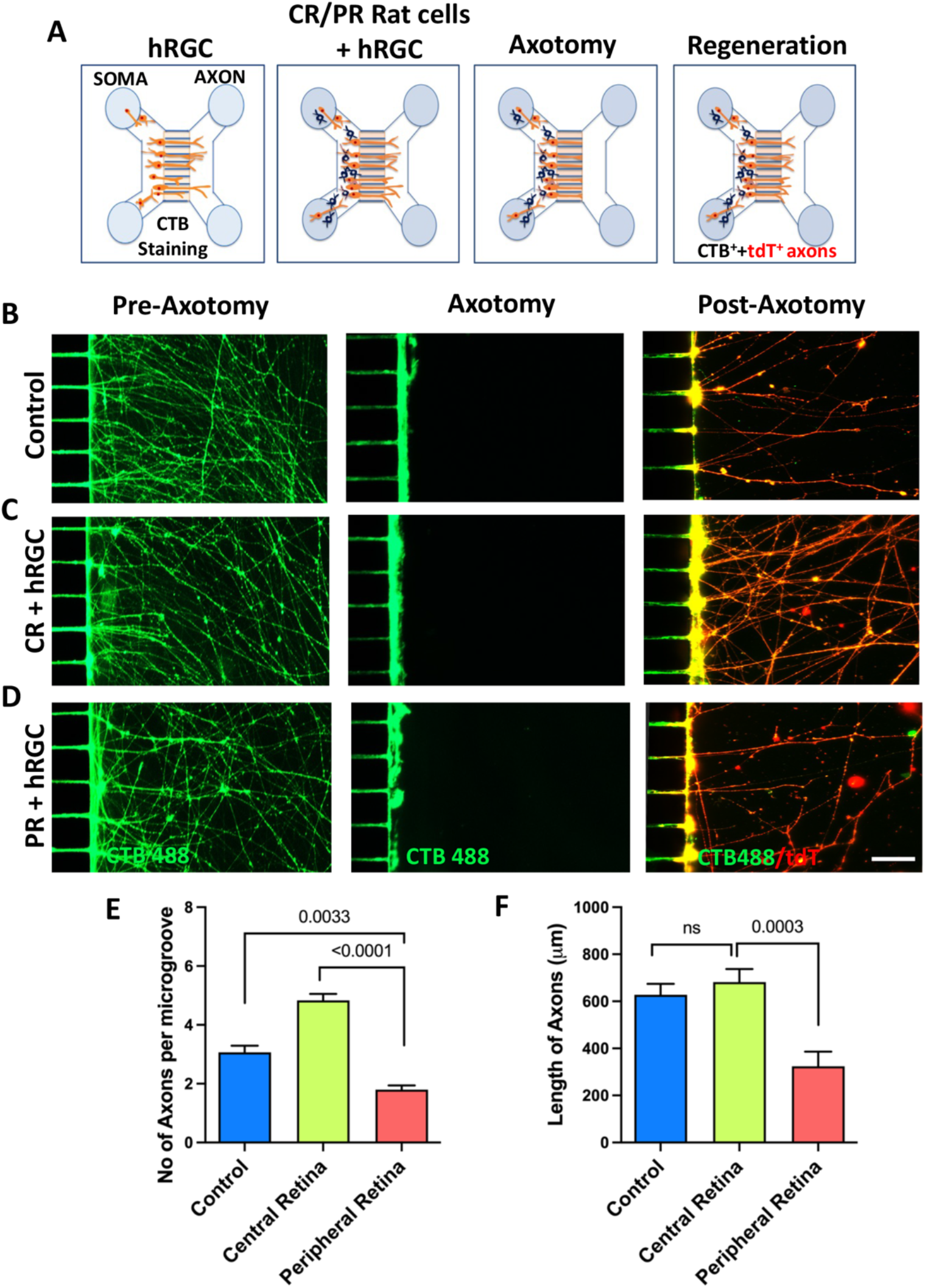
The impact of spatially arrayed chemotropic on hRGCs axon regeneration: **A**. hRGCs^Brn3b-tDT^ were seeded in the soma chamber of a microfluidic device till the axons filled all microgrooves and reached axon chamber. Central/peripheral E16 rat retinal cells were introduced in the soma chamber to co-culture them with already seeded hRGCs^Brn3b-tDT^. After staining with CBT 488 axotomy was performed followed by quantification of regenerated axons (tdT^+^ CBT^+^). **B-F**. hRGCs^Brn3b-tDT^ co-cultured with central rat retinal cells regenerated more axons (C, E) than those cultured alone (B, E) or co-cultured with peripheral rat retinal cells (D, E). The length of regenerated axons was significantly long in hRGCs^Brn3b-tDT^ co-cultured with central (C, F) versus peripheral (D, F) rat retinal cells. There was no significant difference in axon length between control and central retinal groups (B, C, F). Values are expressed as mean±s.e.m. (Student’s t-test; P<0.05, ns, not significant). Experiments were carried out in triplicates per group. Scale bars: 50 μm.

**Fig. 10.**
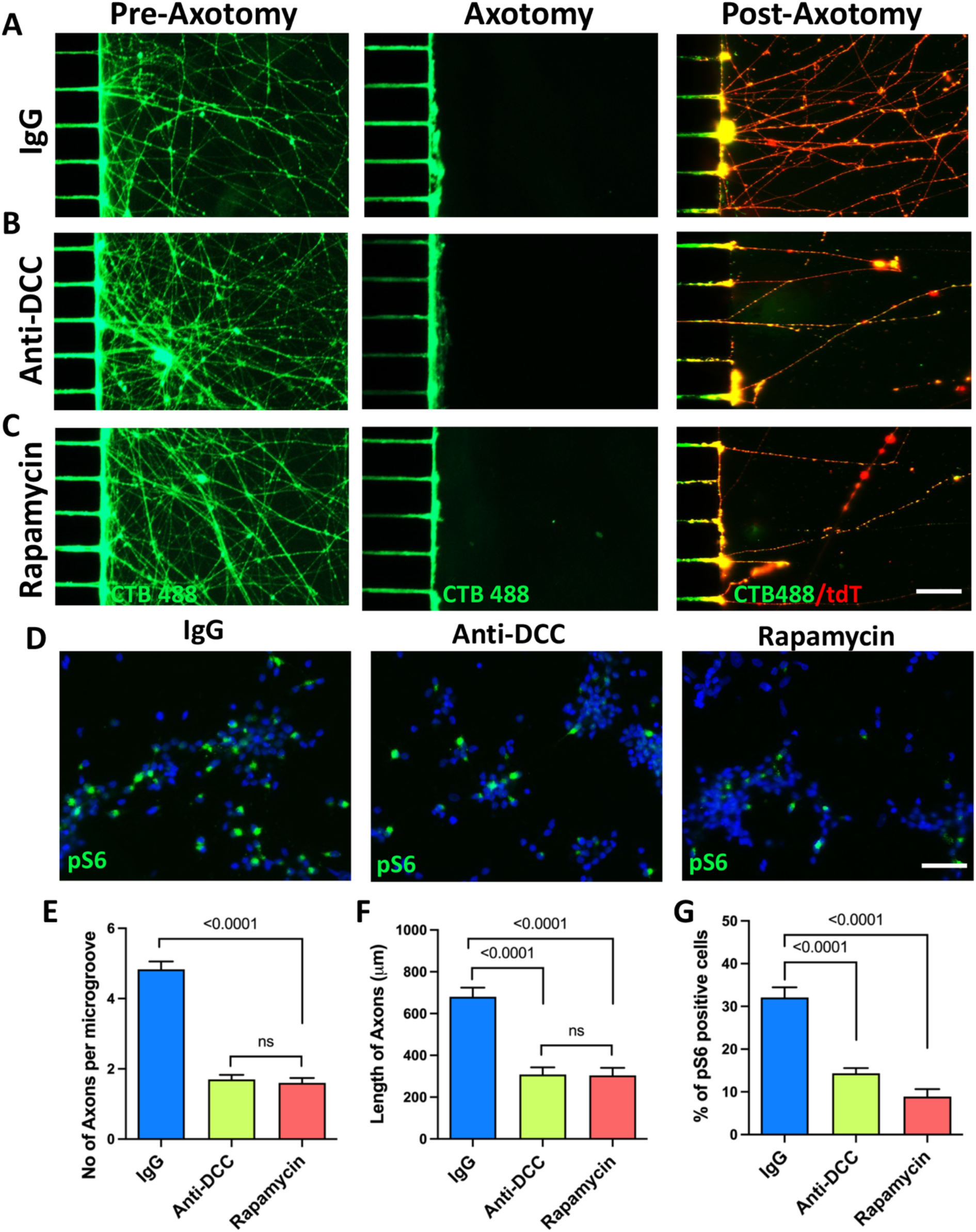
The influence of Netrin-1/DCC interactions and mTOR pathway on hRGCs axon regeneration: **A-C**. hRGCs^Brn3b-tDT^ were seeded in the soma chamber of a microfluidic device till the axons filled all microgrooves and reached axon chamber. Central E16 rat retinal cells were introduced in the soma chamber to co-culture them with already seeded hRGCs^Brn3b-tDT^ which were preincubated with IgG control (A) or anti-DCC antibody (B) or treated with rapamycin (C) After staining with CBT 488 axotomy was performed followed by quantification of regenerated axons (tdT^+^ CBT^+^). **D**. Immunoreactivities corresponding to pS6, as a readout of the mTOR signaling was determined by immunofluorescence analysis. **E-G**. Both the number and length of regenerated axons were reduced in anti-DCC antibody (B, E, F) and rapamycin (C, E, F) groups versus controls (A, E, F). The proportion of hRGCs with immunoreactivities corresponding to pS6 was significantly reduced in anti-DCC antibody and rapamycin groups versus IgG controls (D, G). Values are expressed as mean±s.e.m. (Student’s t-test; P<0.05, ns, not significant). Experiments were carried out in triplicates per group. Scale bars: 50 μm.

### Chemotropic influence on hRGCs is reflected in Netrin-1/DCC recruited transcriptomes

The chemotropic cues, besides facilitating axon guidance, influence diverse biological processes (Lai Wing Sun et al., 2011, Glasgow et al., 2018, Blockus and Chedotal, 2016, Gonda et al., 2020), presumably by recruiting multiple intracellular pathways (Boyer and Gupton, 2018). To identify genes and pathways recruited by Netrin-1/DCC interactions for neuritogenesis, neural activity, and regeneration, we examined the transcriptional profile of hRGC ^Brn3b-tdT^ co-cultured with the central E16 rat retinal cells with or without anti-DCC antibody. The experiment was carried out as described in Figure 3 except that the hRGC^Brn3b-tdT^ were grown on glass coverslips before being co-cultured inverted on the central/peripheral E16 rat retinal cells to retrieve hRGCs uncontaminated with rat cells for RNA seq analysis (Teotia et al., 2017b)(**Fig. 11**). Differentially expressed genes (DEGs) (Love et al., 2014, Team, 2013) were first assigned to GO terms to determine their association with specific biological processes underlying Netrin-1/DCC mediated influence on hRGC axon growth and function. Among the top ranking GO terms “Membrane Potential” and “Axon Guidance” were selected for the hierarchical cluster analysis. The GO term, “Regeneration” contained majority of genes that were exclusively inflammation related and therefore reflected the stress-response of hRGCs to the blocking of Netrin-1/DCC interactions, which is a topic of a separate publication (Subramani and Ahmad). The The genes that were down regulated in response to blocking of Netrin-1/DCC interactions were examined as potential mechanistic targets. For example, in the “Membrane Potential” GO term the majority of genes that may influence membrane potential-exemplified by voltage gated Na channels (e.g., *SCN3A, SCN3B, SCN2A, SCN8A, SCN7A*), voltage-gated calcium channels (e.g., *CACNG2, CACNB2*), voltage-gated potassium channels (e.g., *KNNA2, KCNH4, KCNH8, KCND2, KCNB1,KCNIP1*)-were downregulated in response to the blocking of Netrin-1/DCC interactions suggesting their involvement in Netrin-1 mediated neural activity observed in hRGCs^Brn3b-tdT^ (**Fig. 11C**). In the “Axon Guidance” GO term, among the downregulated genes in response to blocking of Netrin-1/DCC interactions included *DCC* along with other Netrin-1 receptors, *UNC5C* and *UNC5D*, co-expressed in many neurons including RGCs (**Fig. 11D**)(Lai Wing Sun et al., 2011, Murcia-Belmonte et al., 2019). This result is informative as positive influence of Netrin-1/DCC interactions on Netrin-1 receptors gene expression may offer a mechanistic explanation for the diverse cellular effects of Netrin-1 on human RGCs. For example, the binding of Netrin-1 with DCC leads to the recruitment of diverse intracellular pathways including those that mediate protein translation (extracellular kinase 1 and 2 (ERK1, 2); phosphatidylinositol-3Kinase-AKT pathway-mTOR pathway), cytoskeletal rearrangements (actin-binding proteins, Enabled/vasodilator-stimulated phosphoprotein ENA/VASP and neuronal Wiskott-Aldritch syndrome protein; N-WASP), intracellular calcium mobilization (activation of protein kinase C (PKC))(Lai Wing Sun et al., 2011), and membrane potential (Src-mediated activation of NMDAR) (Horn et al., 2013, Glasgow et al., 2018). That some of these diverse pathways were recruited by Netrin-1/DCC interactions in hRGCs^Brn3b-tdT^ was demonstrated by mapping the DEGs on the KEGG pathways (**Fig. 12**), which identified the influence of blocking Netrin-DCC interactions on higher level system functions. The analysis revealed that the majority of genes in the MAPK pathway (**Fig. 12B**) and PI3K-AKT pathway (**Fig. 12C**) were inhibited when Netrin-1/DCC interactions were blocked, suggesting the involvement of these pathways in axon growth/guidance and synaptogenesis, as observed previously. For example, Netrin-1 mediated modulation of protein translation and degradation through the recruitment of the PI3K-AKT pathway in the growth cones constitute an important mechanism for axon guidance (Campbell and Holt, 2001). Similarly, Netrin-1/DCC mediated activation of MAPK was shown to be required for axon growth and orientation (Forcet et al., 2002). Besides axon guidance and growth, Netrin-1/DCC interactions facilitate synaptogenesis by recruiting the mTOR pathway, a downstream hub of the PI3K-AKT pathway, for local protein syntheses in the maturing synaptic structure (Goldman et al., 2013). Our observation that Netrin1/DCC interaction-mediated axon regeneration was rapamycin-sensitive suggested the involvement of the mTOR pathway, corroborated by the transcriptional recruitment of the PI3K-AKT pathway. Of the rest, the Hippo signaling (**Fig. 12D**) and p53 pathways (**Fig. 12E**) were of especial interest as they could affect the diverse functions influenced by Netrin-1/DCC interactions. For example, the Hippo pathway, which regulates gene expression through the homologous transcription co-activator YAP/TAZ and the TEAD transcription factor, does not have a designated ligand or receptor but integrates multiple cellular and extra-cellular signals through cross talk with other signaling pathways, including the PI3K pathway (Moya and Halder, 2016). Therefore, the recruitment of the Hippo pathway by Netrin-1/DCC interactions may add to the versatility of Netrin-1 signaling. The recruitment p53 signaling may further add to that versatility as p53 could transcriptionally regulate the expression of Netrin-1 and its receptors (Arakawa, 2005), thereby affecting Netrin-1 and DCC interactions and hence Netrin-mediated diverse functions.

**Fig. 11.**
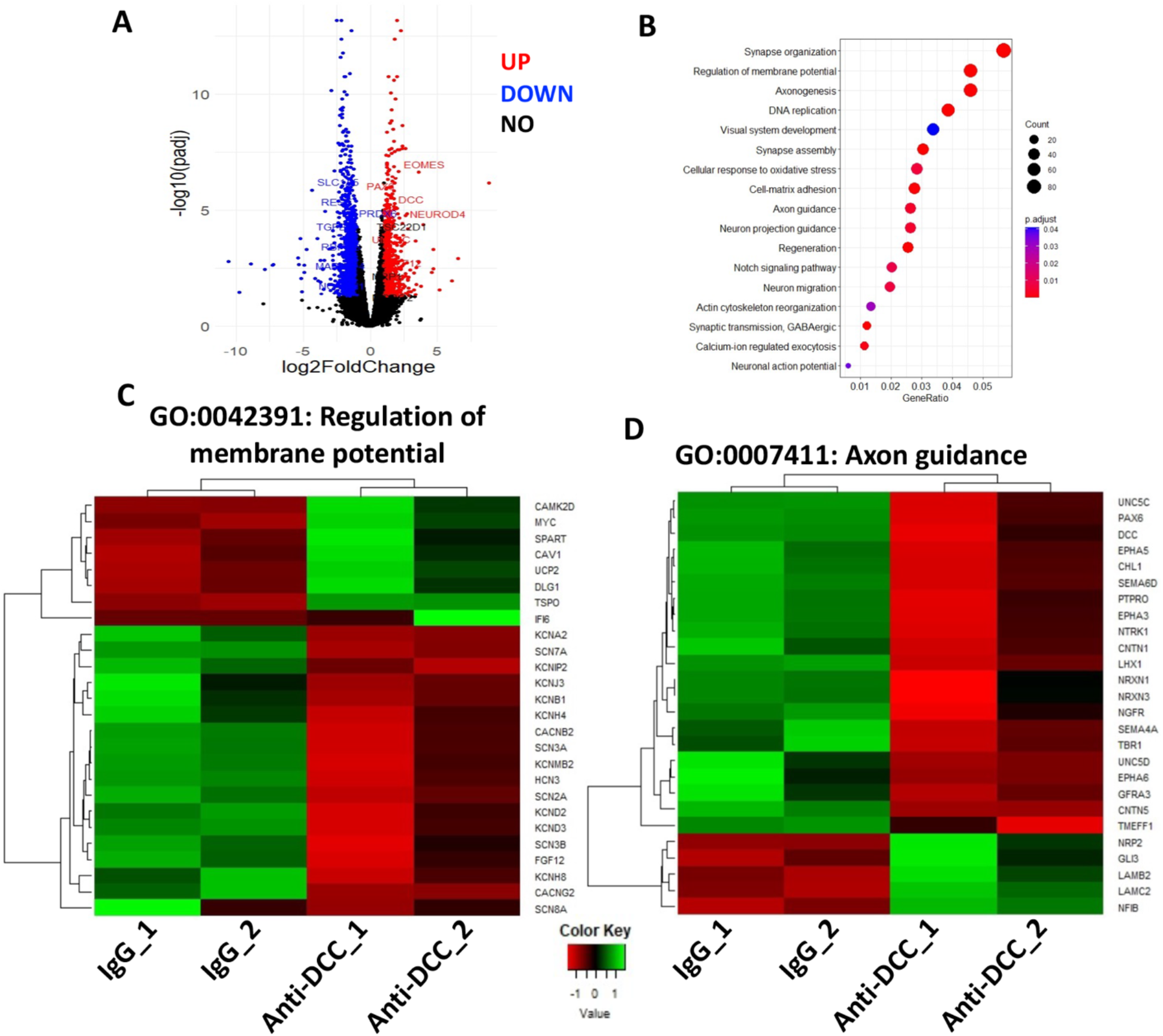
Impact of Netrin-1/DCC interactions on hRGCs’ transcriptome: RNAseq analysis was carried out on hRGCs^Brn3b-tdT^, co-cultured with central E16 rat retinal cells, pre-exposed to IgG or anti-DCC antibody to determine the effects of Netrin-1/DCC interactions on global gene expression. **A**. Representative volcano plot of differentially expressed genes (padj-value < 0.05, two-fold change) in IgG-vs Anti-DCC antibody-treated groups. **B**. Significantly enriched gene ontology (GO) terms are represented for DEGs in hRGCs between IgG and Anti-DCC antibody treated groups in a bubble plot. **C, D**. Heat maps and hierarchical clustering-based dendrograms of DEGs between IgG control and Anti-DCC groups show that the global gene expression pattern differed in select GO terms such as the “regulation of membrane potential” (C) and “Axon guidance” (D). Inhibition of Netrin-1/DCC interactions by anti-DCC antibody decreased the expression of genes belonging to the above-mentioned GO terms compared to IgG controls. Expression variability between samples was indicated by Z score, where shades of red and green color denote down and upregulation of genes, respectively.

**Fig. 12.**
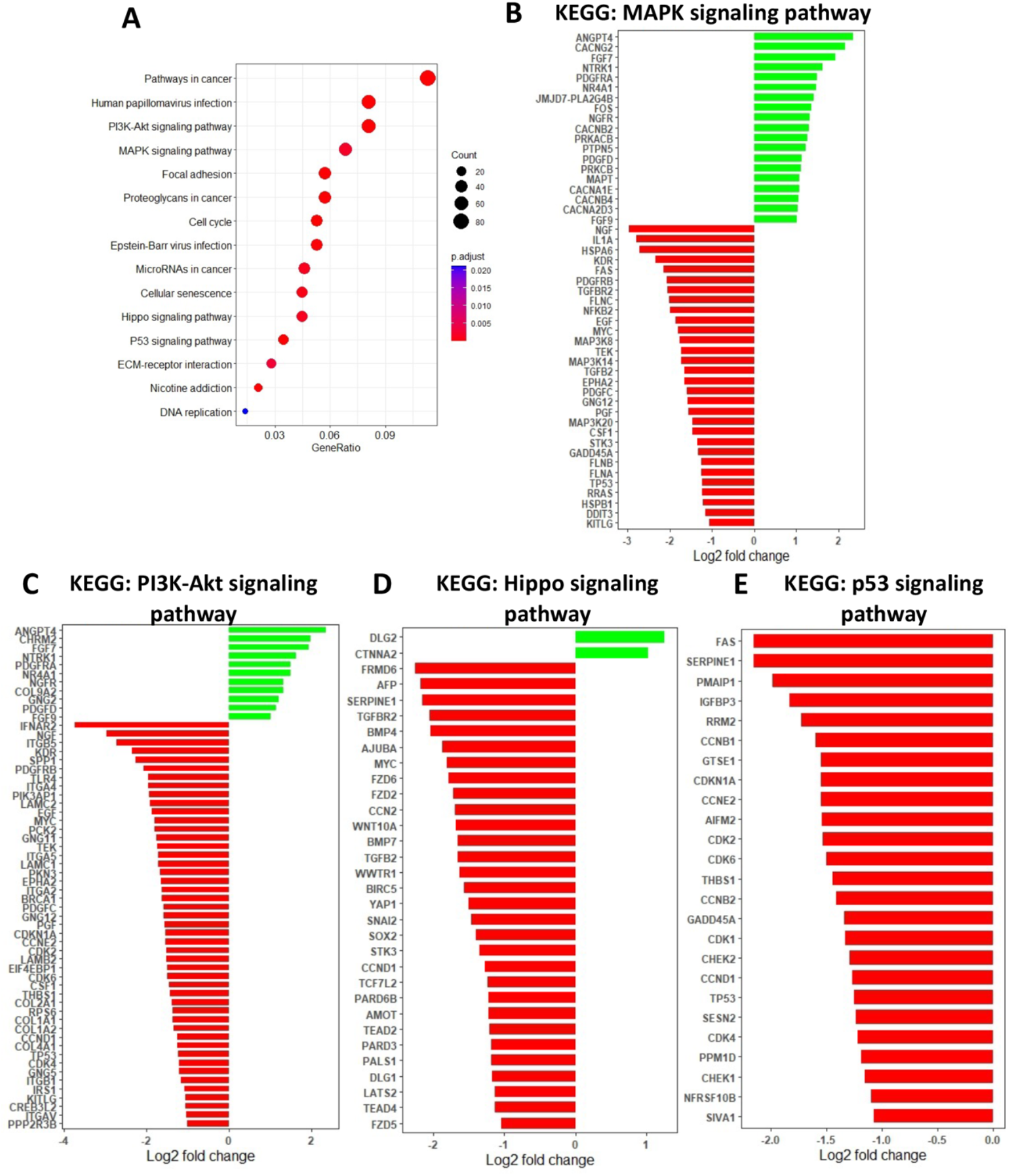
Impact of Netrin-1/DCC interactions on hRGCs’ genome pathways: **A**. Kyoto Encyclopedia of Genes and Genomes (KEGG) pathway analysis mapped DEGs between the IgG controls and anti-DCC antibody group on the top 15 KEGG pathways as displayed in the bubble plot. **B-E**. Representations of DEGs (log2 fold change) mapped on MAPK pathway (B), PI3K-Akt signaling pathway(C), Hippo Signaling pathway (D), and P53 pathway (E) are displayed and MAPK signaling pathway (D) are given. These pathways are suppressed when Netrin-1/DCC interactions are blocked by anti-DCC antibody, positing them as relevant in mediating Netrin-1 influence on hRGCs functions that include axon growth, neural activities, and regeneration.

## DISCUSSION

The projection of RGC axons to central targets is a complex process involving several stages including guidance within the retina toward the optic disc, decussation (or not) at the optic chiasm, and formation of retinotopic connections at SC and LGN (Erskine and Herrera, 2014, Harada et al., 2007, Oster and Sretavan, 2003). This process is facilitated by an array of evolutionarily conserved guidance molecules, recognized by their cognate receptors on the growth cones (GC) of RGC axons. The information on the mechanism underlying the guidance of RGC axons has largely emerged from studies in rodents and lower vertebrates (Oster and Sretavan, 2003) and whether or not hRGCs recruit similar mechanisms remains poorly understood, primarily due to inaccessibility of hRGCs due to their fetal generation and the fact that their wiring is largely completed by birth (Dreher and Robinson, 1991). Using hRGCs directly generated from pluripotent cells by recapitulation of developmental mechanisms and examining their specific response to guidance molecules arrayed in the developing rat retina, we have begun to study the axon guidance potential of hRGCs and its underlying mechanism. Here, we have demonstrated that developing hRGCs may recruit a similar axon guidance mechanism as observed in the developing rodent retina (Brittis and Silver, 1995, Harada et al., 2007). This notion is supported by the observation that hRGCs recognize and respond to spatially arrayed intra-retinal guidance cues for the centripetal polarization of axons, necessary for their exit at the optic disc, the first stage in the navigation toward the central targets. We observe that the central retinal cells can mediate significant chemoattraction through Netrin1-/DCC interactions and regulate central orientation of hRGCs axons toward the optic disc. This observation appears contradictory to previous finding that Netrin-1 mediated chemoattraction is needed for facilitating hRGC axons’ entrance into the optic disc and not for their central projection because in the Netrin-1 KO mice RGC axons reach the optic disc but fail to enter the optic nerve (Deiner et al., 1997). However, the presence of directionless axons in the central retina in the Netrin-1 KO mice (Deiner et al., 1997) suggests that Netrin-1/DCC interaction in the central retina, observed here and supported by Netrin-1 immunoreactivities in the central retinal cells, may maintain the central trajectory of RGC axons once it is set by the chemo-repulsive environment in the peripheral retina.

In the developing retina, cells are born in a central-to-peripheral gradient such that at a given embryonic stage, for example at E16, cells in the central retina are relatively more developmentally mature than the peripheral retina (Rapaport et al., 2004, Lo Giudice et al., 2019). However, whether or not the developmental gradient is reflected in the neural activity of the central versus peripheral RGCs and if it is regulated by spatial extrinsic cues is not known. Using hRGCs in the co-culture paradigm as surrogates for the central versus peripheral RGCs, we observed that these cells may differ in physiological paradigms, which are significantly influenced by spatially arrayed Netrin-1/DCC, Slit/ROBO2, and CSPG interactions. The observation predicts that in the developing human retina, as in the rodent retina, spatially arrayed chemotropic cues may contribute toward the neuronal activities of RGCs besides regulating neuritogenesis and axon growth. Given the facilitatory role of neuronal activity on Netrin-mediated axon growth (Bouchard et al., 2008) and the recent evidence Netrin-1/DCC interactions facilitate on LTP (Horn et al., 2013) suggest that these activities act synergistically for interneural connections and communication. These two processes, facilitated by Netrin-1/DCC interactions, may also underlie better regeneration capacity of hRGCs axons, when co-cultured with the E16 central retinal cells.

Besides, the regeneration may also be aided by mTOR signaling as the inhibition of the mTOR complex by rapamycin abrogated Netrin-1 mediated improvement in regeneration. Netrin-1/DCC interactions are known to recruit several intracellular pathways that include phosphoinositide 3 kinase (PI3K)/Akt and mitogen activated protein kinase (MAPK) (Boyer and Gupton, 2018), activated via the focal adhesion kinase (FAK), the downstream mediator constitutively bound to DCC, and both PI3K/Akt and MAPK pathway can cross talk to regulate mTOR signaling (Mendoza et al., 2011). mTOR signaling has been demonstrated to facilitate regeneration in an optic nerve crush model (Park et al., 2008) and of hRGC axons following chemical axotomy (Teotia et al., 2019). Netrin-1 mediated regeneration of the optic nerve has been observed in lower vertebrates like goldfish (Petrausch et al., 2000). Whether or not that involves recruitment of the mTOR pathway remains unknown. However, the observations that Netrin-1/DCC interactions in adult CNS appear to inhibit axon regeneration (Ellezam et al., 2001, Petrausch et al., 2000, Low et al., 2008, Manitt et al., 2004) suggest that netrin-mediated regeneration observed in lower vertebrates is confined to the developmental stage in higher vertebrates. It is not known why RGC axon regeneration becomes refractory to Netrin-1/DCC interactions in adult. That Netrin-1/DCC interactions in adult RGC fail to recruit the mTOR signaling can be a possibility. In summary, our results demonstrate that during development, human RGCs may follow an evolutionarily conserved mechanism of intra-retinal guidance to locate and exit at the optic disc, the first step toward connecting with the central targets. This mechanism may include Netrin-1/DCC - mediated neural activity and axon growth, presumably acting synergistically. This may underlie Netrin-1/DCC-mediated axon regeneration aided by the recruitment of mTOR signaling to facilitate local protein synthesis for axon growth. Besides shedding light on the development and guidance of hRGC axons, our findings demonstrate that human pluripotent cell derived RGCs may facilitate stem cell approach to glaucomatous degeneration where success requires faithful navigation of axons to appropriate central targets.

## MATERIALS AND METHODS

### Experimental animals

The use of animals and experimental protocols were approved by the Institutional Animal Care and Use Committee at the University of Nebraska Medical Center (UNMC) and conducted in accordance with the Association for Research in Vision and Ophthalmology (ARVO) Statement for the Use of Animals in Ophthalmic and Vision Research. Postnatal day 15 (PN15) and embryonic day 16 (E16) timed-pregnant Long Evans rats (Charles River Laboratories) were used for harvesting retinal explants/cells from P15 and E16 embryos retina. Rats were euthanized by CO2 exposure followed by decapitation. Retinae were dissociated using the Papain Digestion Kit (Worthington Biochemical Corporation, Lakewood, NJ) to obtain single cells as per vendor instructions.

### Directed differentiation of hES cells into RGCs

Differentiation of Brn3b reporter hESCs^Brn3b-tdT^ (Sluch et al., 2017) into functional RGCs was achieved through a modified stage-specific chemically defined protocol for directed differentiation of pluripotent cells into RGCs (Teotia et al., 2017a). Briefly, to enhance the reproducibility of RGC generation NRs were generated in high density monolayer culture of ES cells on 1% Matrigel (Corning) in the presence of dual SMAD inhibitors [LDN (100nM) and SB325 (10uM)] for the first three days. The medium was changed every day, and the culture was continued for the next 6 days in the presence of DKK (10ng/ml), IGF1 (10ng/ml), and LDN (100nM) till NRs with RPC (Rx^+^ Pax^+^ cells) appear. The basal medium for RPC generation contained 2% B27 supplement (Gibco) and 1% N2 supplement in DMEM: F12 (Gibco). The RPCs were plated onto Matrigel-coated dishes (Corning), and RGC differentiation was initiated by treating cells for 2 days with Shh (250ng/ml), FGF8 (100ng/ml) and DAPT (3μM). RGC differentiation was facilitated by treatment with follistatin (100ng/ml), Shh (250ng/ml) and DAPT (3 μM) for 3 days. Finally, RGC maturation and survival, characterized by (*Brn3b*-tdT^+^ Pax6^+^ cells) were promoted by supplementing the medium with BDNF (100ng/ml), forskolin (10μM), NT4 (5ng/ml), CNTF (10ng/ml), cAMP (100μM), Y27632 (1 μM) and DAPT (3μM) for the next 10 days. The medium (RGC maturation medium) was changed every day. The basal medium for RGC generation contained 50% DMED/F12, 50% Neuro Basal media, 0.05% N2 supplement (Gibco), 1% B27 supplement, 5% L-glutamine, 1% β-Mercaptoethanol, BSA (1μg/ml), Insulin (5μg/ml), Sodium Selenite (3nM), AND Apo-Transferrin (50μg/ml). All growth factors and small molecules were purchased from R&D Systems, Sigma and Milteny Biotech.

### hRGCs and E16 rat retinal cells/explants co-culture

The papain-dissociated E16 central/peripheral rat retinal cells were seeded onto Matrigel-coated dishes (Corning) at a density of 1×10^6^ cells in RGC maturation medium. hRGCs ^Brn3b-tdT^ were seeded on the rat retinal cells at the density of 1×10^5^ cells and cultured for 5 days with daily medium change. hRGCs ^Brn3b-tdT^ were treated with human IgG or anti-DCC antibody (10μg/ml; BD Biosciences) or anti-Robo2 antibody (20μg/ml; Santa Cruz) before co-culture. In some instance E16 peripheral rat retinal cells were treated with chondroitinase ABC (0.2 U/ml; Sigma-Aldrich) to degrade CSPG before culture. After 5 days *in vitro* (DIV), hRGCs ^Brn3b-tdT^ were subjected to Sholl and electrophysiological analyses. The E16 rat retinal explants were cultured on inserts (pore size, culture plate, vendor) at air/medium interface. IgG-/anti DCC antibody-treated hRGCs ^Brn3b-tdT^ 1×10^5^ cells were seeded on the explants and cultured for 5 days, following which they were subjected to two-photon microscopy in live culture or fixed in 4% paraformaldehyde and frozen for immunofluorescence analysis, carried out on the cryosection.

### Chemical axotomy and Regeneration

Chemical axotomy and regeneration was carried out as previously described (Teotia et al., 2019). Briefly, Polydimethylsiloxane microfluidic devices with 450 μm microgrooves (SND 450, Xona Microfluidics) were assembled and prepared as per the manufacturer’s instruction with the cover glass coated overnight with 500μg/ml poly-D-lysine (Sigma-Aldrich). Once assembled, a solution of 1% Matrigel (Corning) in DMEM/F-12 medium (Invitrogen) was added to the four reservoirs/well (both side soma and axonal compartment) of the device. Devices were coated with Matrigel for at least 1 hour at room temperature before cell seeding. hRGCs ^Brn3b-tdT^ were re-suspended at 2×10^4^ cells/μl in RGC maturation and survival media and loaded into the micro-channel in a 5μl droplet. Cells were allowed to attach for at least 30 min, after which medium was added into the adjacent wells. The axonal compartment was filled with similar medium to facilitate axonal growth. hRGCs ^Brn3b-tdT^ were cultured for 3 days till all the microgrooves were occupied by the hRGCs ^Brn3b-tdT^ axons. The axons were subjected to retrograde labeling, using 1% cholera toxin subunit B (CTB) Alexa Fluro™ 488 conjugate (Invitrogen) dissolved in RGC media and added to the axonal compartment (100 μl per well) of the microfluidic device and incubated overnight at 37°C. The soma chambers of the devices were seeded with the E16 central/peripheral rat retinal cells 1×10^5^ cells in 5μl were seeded prior to axotomy. Axotomy was performed by first removing media from the axonal compartment and adding 50μl RGC medium with 0.5%saponin (Sigma) for 3 min. To prevent the flow of the detergent into the soma compartment, a hydrostatic pressure was maintained by volume difference between soma (200μl/well) and axonal (50μl/well) compartments. At the end of a 3 min period, followed by two PBS washes, the axonal compartment was re-coated with 1% Matrigel for 30 min at 37°C. After Matrigel coating, media was returned immediately to the axonal compartment for the duration of the culture time. hRGCs ^Brn3b-tdT^, post-axotomy, were cultured for an additional 5 DIV for axonal regeneration. In some groups hRGCs ^Brn3b-tdT^ were pre-treated with anti-DCC antibody to disrupt Netrin-DCC signaling or rapamycin (Cell signaling; 100nM) was added to the medium to inhibit mTOR signaling.

### Immunofluorescence, image acquisition, and data analysis

Immunofluorescence analysis of cells and explants were carried out as previously described (Teotia et al., 2019). Briefly, fixative-treated cells and explants, which were frozen and cryosections were exposed to 5% normal goat/donkey serum in PBS for 30 min at room temperature and permeabilized with Triton X-100 (0.4% or 0.2% for nuclear or cytoplasmic staining, respectively), followed by overnight incubation with primary antibody at 4°C. The next day, cells/sections were treated with fluorescence (Cy3/FITC)-tagged secondary antibodies (Life Technologies) diluted in 5% donkey serum and 0.2%/0.4% Triton X-100 solution for 1 hour at room temperature, followed by three washes with PBS. Samples were mounted using VectaShield (Vector Laboratories) and fluorescent images were acquired with Zeiss ApoTome.2 Imager M2 upright microscope (Axiovert 200M) and Axiovision 4.8 software (Carl Zeiss). The percentage of cells expressing specific markers was determined by counting immune-positive cells in randomly chosen five visual fields per coverslip. For each experimental condition, three biological replicates were used to calculate means and standard deviation. A list of antibodies and working dilutions is provided in Table S1. For growth cone analyses, Alexa Fluor 594-phalloidin staining (Cytoskeleton) for F-actin was carried out by incubation at 1:100 dilution in PBS containing 5% normal goat serum for 30 min after fixation and permeabilization, as per manufacturer’s instruction. Neurite complexity and length quantification was done by Sholl analysis using the FIJI Simple Neurite Tracer (ImageJ, National Institutes of Health) plugin (Longair et al., 2011). Sholl analysis was carried out on each traced RGC with a 10um radius step size. The overall RGC radius was calculated as the distance from the soma’s center to the terminus of the longest neurite. The percentage of tdT^+^FITC^+^ regenerating axons was calculated by counting the numbers of both tdT^+^ FITC^+^ axons out of total numbers of tdT^+^ axons that crossed the microgroove and were present in an axonal compartment at the time of imaging.

### Electrophysiological analysis

Coverslips were affixed to a recording chamber on the stage of an Olympus BX51-WI microscope using vacuum grease and superfused with Ames’ medium (US Biologicals) bubbled with 95% O_2_/5% CO_2_ at room temperature. Cells were targeted for whole-cell patch-clamp recording with pipettes pulled from thin-walled borosilicate glass capillary tubes (1.2 mm OD, 0.9 mm ID) filled with a solution comprised of (in mM) 120 potassium-gluconate, 8 KCl, 2 EGTA, 10 HEPES, 5 ATP-Mg, 0.5 GTP-Na_2_, 5 phosphocreatine-Na_2_, 0.1 lucifer yellow-Li. Pipettes had resistances of 5-8 MΩ. Cells were voltage-clamped at -84 mV (after correction for 14 mV liquid junction potential) and voltage-gated Na^+^ and K^+^ currents were recorded in response to a series of depolarizing voltage steps (150 ms, -74 to +56 mV, 10 mV increments). Series resistance was partially compensated (65-75%). Spiking activity was recorded in response to a series of depolarizing current injections (+2.5 to +15 pA, 2.5 pA increments, 500 ms). Current and voltage-clamp stimuli were controlled with a Multiclamp 700B amplifier and digitized with a Digidata 1550B. For analysis, Na^+^ and K^+^ currents were normalized to cell capacitance measured using the amplifier circuitry and I_Na_ was measured at the peak of the current while I_K_ was measured at the end of the 150 ms stimulus step.

### Quantitative polymerase chain reaction analysis

Quantitative polymerase chain reaction (Q-PCR) analysis was carried out as previously described (Teotia et al., 2019). Briefly, total RNA from cells was extracted using Mini-RNeasy kit (Qiagen) according to the manufacturer’s instructions. One μg of total RNA per sample was used for reverse transcription into cDNA, using Superscript III RT kit, following the manufacturer’s instructions. Q-PCR was performed using Quantifast SYBR Green Master Mix (Qiagen) on Rotor Gene 6000 (Corbett Robotics). All qPCR results represent each sample measured in triplicate. No-template blanks were used for negative controls. Amplification curves and gene expressions were normalized to the housekeeping gene GAPDH. The primer sequences used in this study are listed in Table S2.

### RNA-seq Analysis

RNA-seq analysis was carried out as previously described (Teotia et al 2017). Briefly, 50ng of RNA was utilized to create sequencing libraries for 75bp single end reads using the True Seq mRNA V2 Kit from Illumina, Inc. in San Diego, California. RNA seq data, imported into BaseSpace (illumine, Inc.) for analysis after being recorded in the FASTQ format, were quantified in counts by RSEM. Gene differential expression analyses were conducted with the DESeq2 package in R (Love et al., 2014). Genes with 0 or1 count across all samples were removed and the filtered data were normalized by a median of ratios approach and dispersion was estimated. Negative binomial generalized linear models were used to test differential expression between groups. The Benjamini-Hochberg method was used to control the false discovery rate (FDR) to be no more than 0.05 (Benjamini and Hochberg, 1995). Genes with FDR below 0.05 and at least 2-fold changes were selected as differentially expressed. Independent filtering on the mean of normalized counts was applied to the analysis results to optimize the number of FDR-adjusted p values lower than a significance level. The FDR-adjusted p values for the genes which do not pass the independent filtering threshold were set to NA. The Kyoto Encyclopedia of Genes and Genomes (KEGG) pathways analysis and Gene ontology were both used to map the DEGs discovered at each step. Gene Ontology Consortium (http://geneontology.org) was utilized for gene ontology enrichment analysis, whereas Profiler was used for KEGG enrichment analysis.

### Statistical analysis

The mean S.E.M. is used to express values. GraphPad Prism was used to analyze and visualize the data (GraphPad). A two-way analysis of variance (ANOVA) or an unpaired Student’s t-test (two-tailed) were used to establish the statistical significance. P-values of 0.05 or lower were regarded as significant. When each group had at least three replicate samples from three different experiments, statistical analysis was done.

## ACKNOWLEDGENTS

Thanks are due to Dr. Don Zack for BRN3b-reporter hESC line. This research was supported by R01EY022051 and R01EY029778 to IA. MJVH is supported by R01EY030507.

## Supplementary Figures

**Fig. S1.**
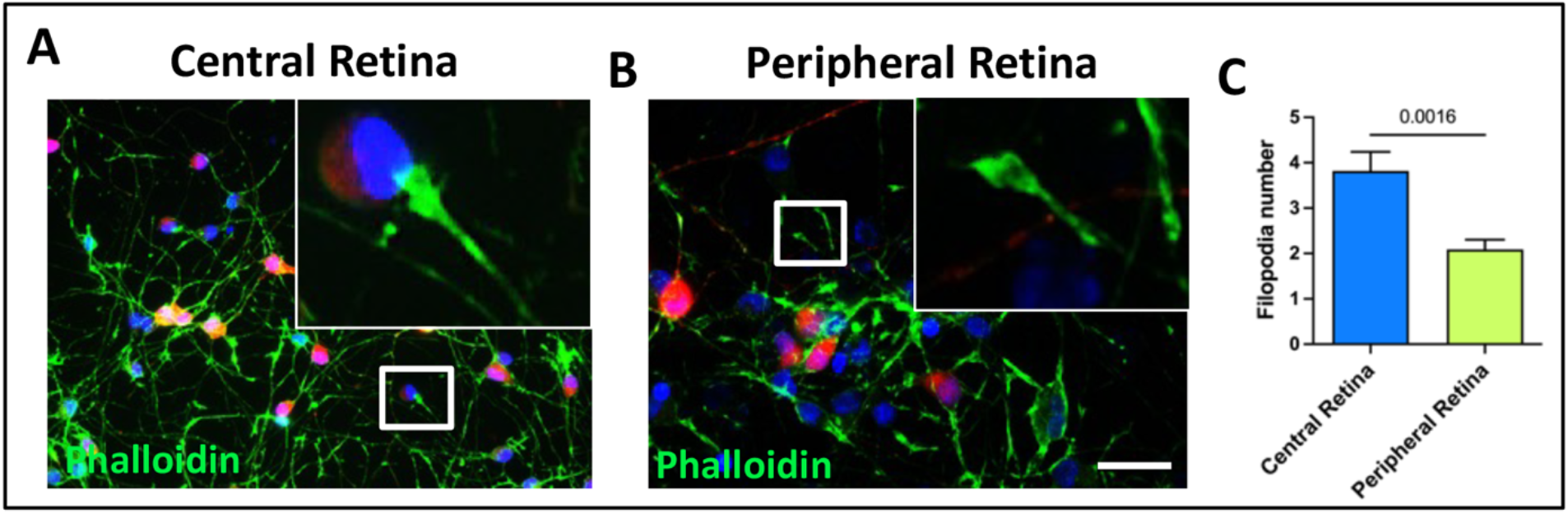
The impact of central versus peripheral rat retinal cells on hRGCs growth cones (GCs): **A-C**. Counting of filopodia in the GCs of hRGCs^Brn3b-tDT^ stained with phalloidin revealed more active GCs when co-cultured on the central versus peripheral rat retinal cells. Values are expressed as mean±s.e.m. (Student’s t-test; P<0.05, ns, not significant). Experiments were carried out in triplicates per group. Scale bars: 50 μm.

**Fig. S2.**
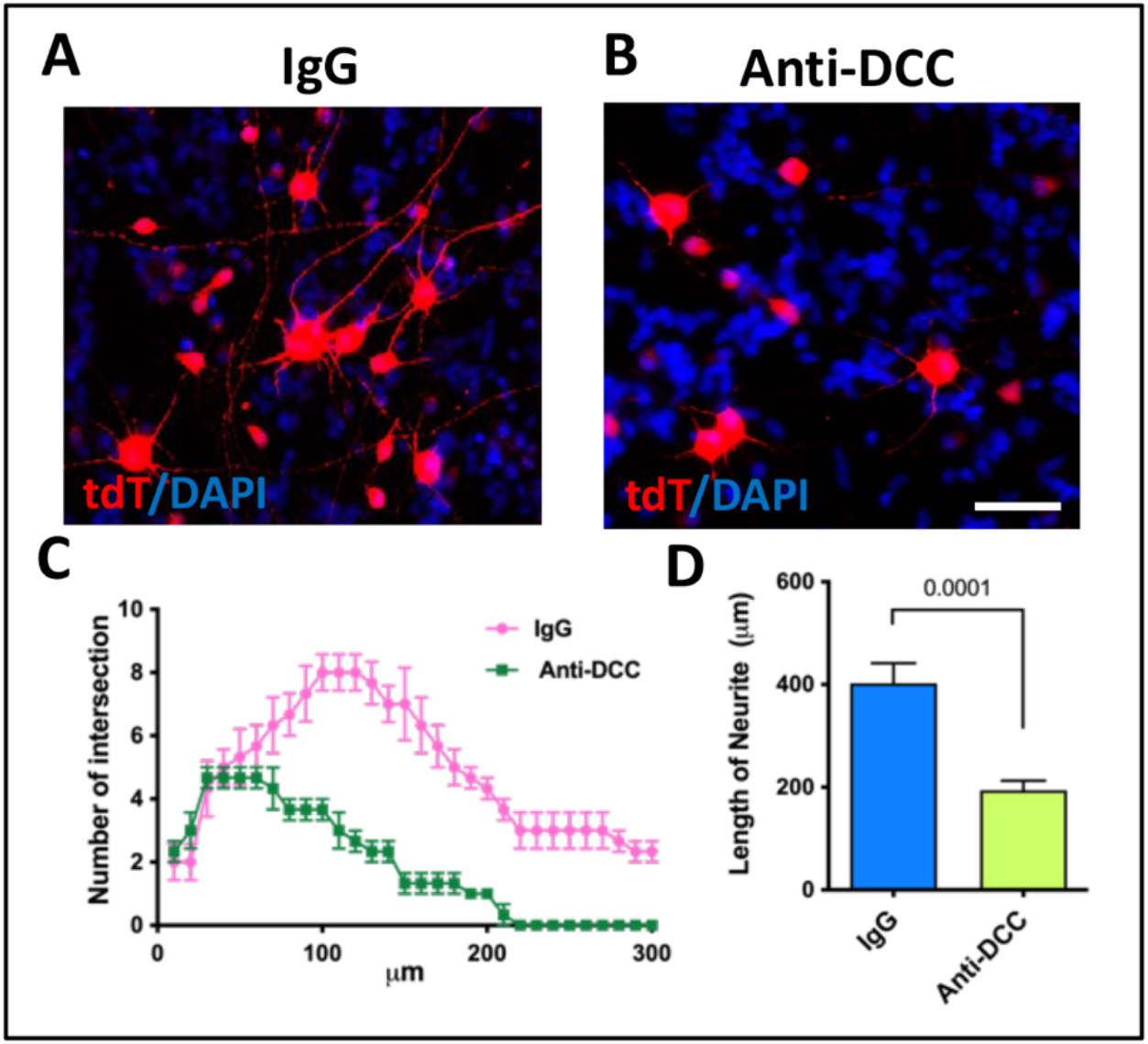
Netrin-1/DCC interactions in mediating chemotropic influence of central rat retinal cells on hRGCs neurites: **A-D**. hRGCs^Brn3b-tDT^ preincubated with IgG (A) and DCC-antibody (B) were co-cultured on E16 central retinal cells followed by Sholl analyses (C, D) of hRGCs^Brn3b-tDT^ neurites. The complexity of axons and their length were significantly decreased when DCC on hRGCs^Brn3b-tDT^ was neutralized by DCC antibody, compared to IgG controls, suggesting Netrin-1/DCC interactions underlying central retinal cells-mediated chemoattraction. Values are expressed as mean±s.e.m. (Student’s t-test; P<0.05, ns, not significant). Experiments were carried out in triplicates per group. Scale bars: 50 μm.

**Fig. S3.**
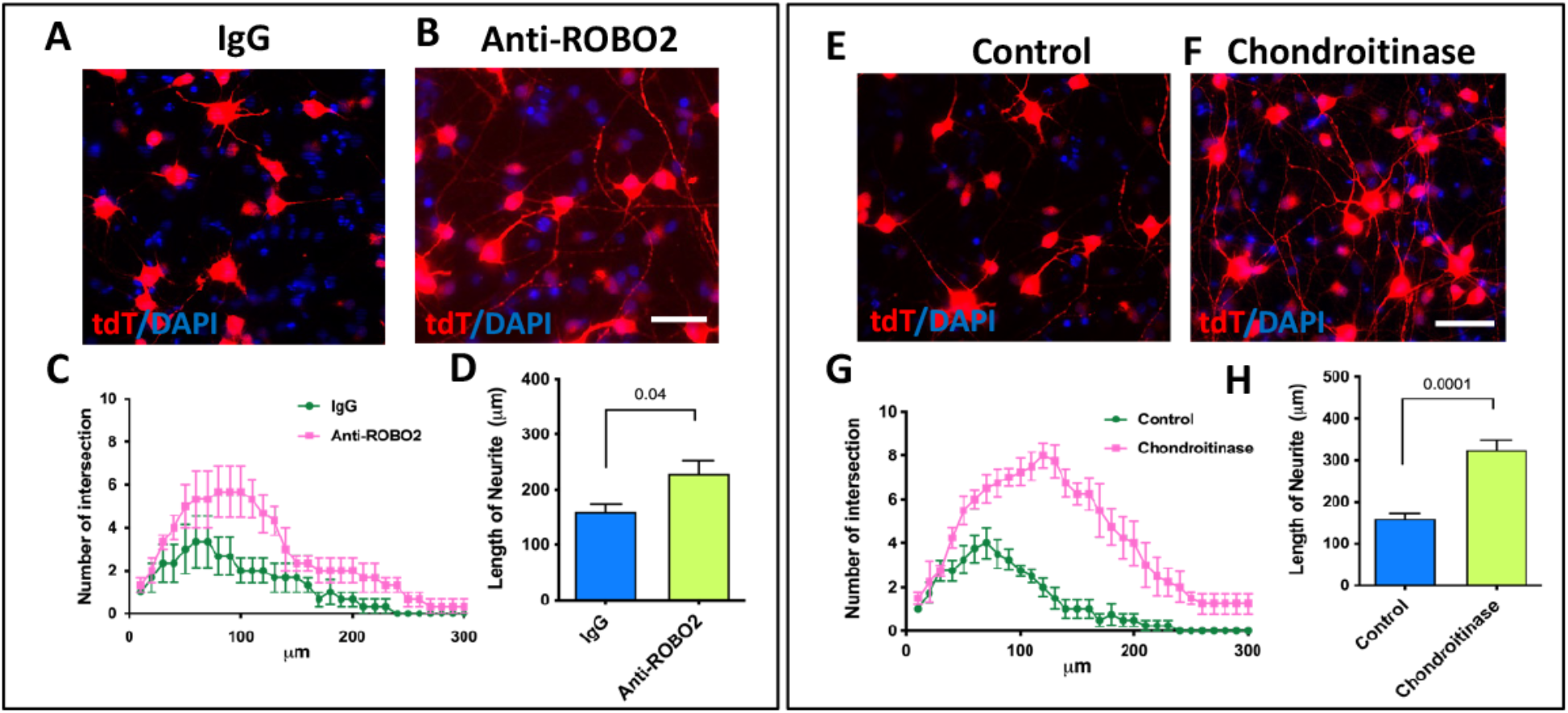
Slit/ROBO2 interactions and CSPG influence in mediating chemotropic influence of the peripheral rat retinal cells on hRGCs neurites: **A-D**. hRGCs^Brn3b-tDT^ preincubated with IgG (A) and ROBO2-antibody (B) were co-cultured on E16 peripheral retinal cells followed by Sholl analyses (C, D) of hRGCs^Brn3b-tDT^ neurites. The complexity of axons and their length were significantly increased when Slits on hRGCs^Brn3b-tDT^ was neutralized by ROBO2 antibody, compared to IgG controls, suggesting Slit/ROBO2 interactions underlying peripheral retinal cells-mediated chemo-repulsion. **E-F**. hRGCs^Brn3b-tDT^ were co-cultured on E16 peripheral retinal cells, untreated (E) and treated with chondroitinase (F) followed by Sholl analyses (G, H) of hRGCs^Brn3b-tDT^ neurites. The complexity of axons and their length were significantly increased when hRGCs^Brn3b-tDT^ were cultured on cells treated with chondroitinase, compared to untreated controls, suggesting that CSPG distributed on peripheral retinal cells underlie peripheral retinal cells-mediated chemo-repulsion. Values are expressed as mean±s.e.m. (Student’s t-test; P<0.05, ns, not significant). Experiments were carried out in triplicates per group. Scale bars: 50 μm.

**Fig. S4.**
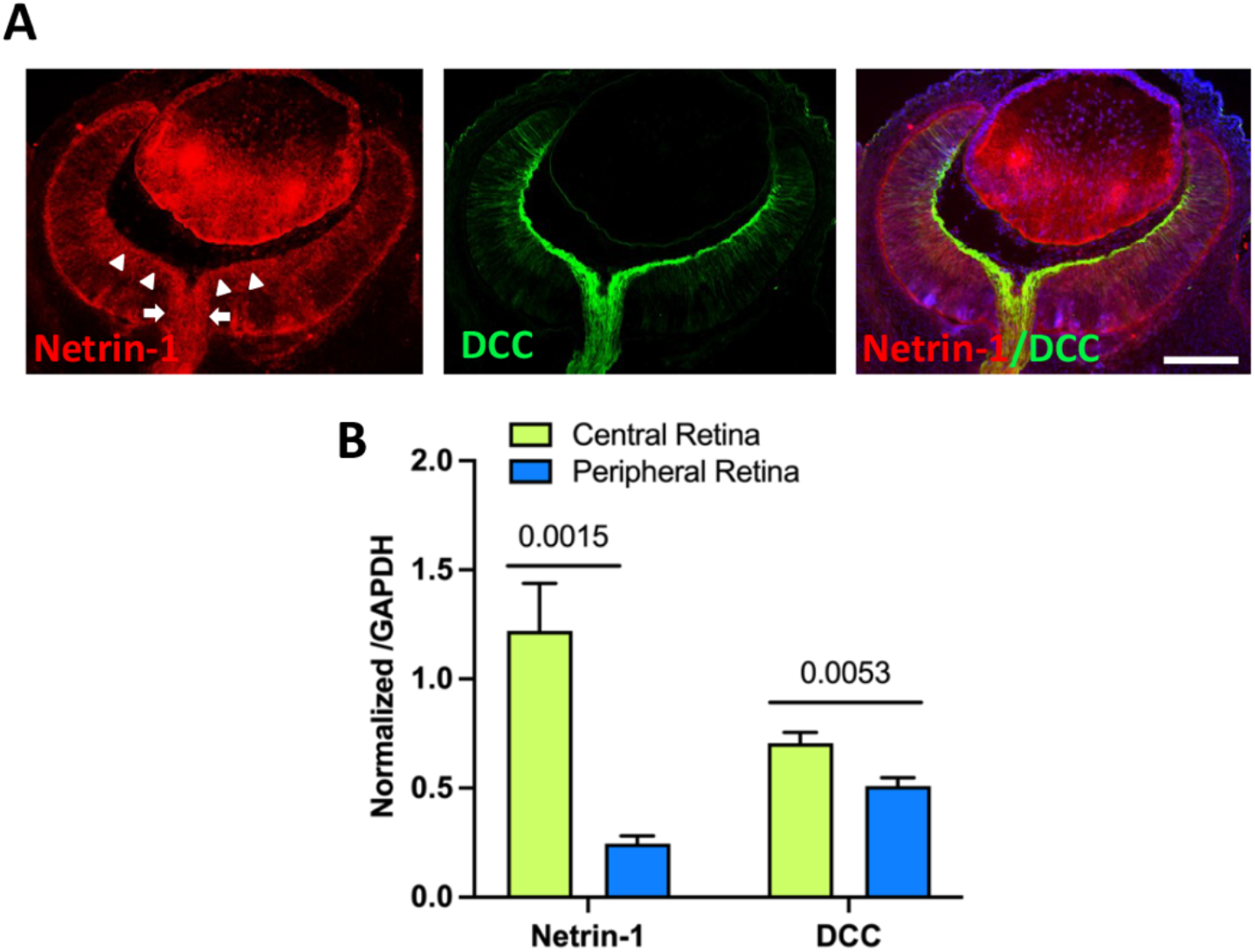
Expression of Netrin-1 immunoreactivities in the developing rat retina: **A**. Spatial distribution of immunoreactivities corresponding to Netrin-1 and ROBO2 in E16 rat retina. Netrin-1 immunoreactivities preferentially localized in the central retina (arrowhead) and optic disc (arrow) as DCC’s. **B**. Spatial distribution of transcripts corresponding to NETRIN-1 and DCC in E16 rat retina as determined by q-PCR analysis. Values are expressed as mean±s.e.m. (Student’s t-test; P<0.05, ns, not significant). Experiments were carried out in triplicates per group. Scale bars: 50 μm.

**Fig. S5.**
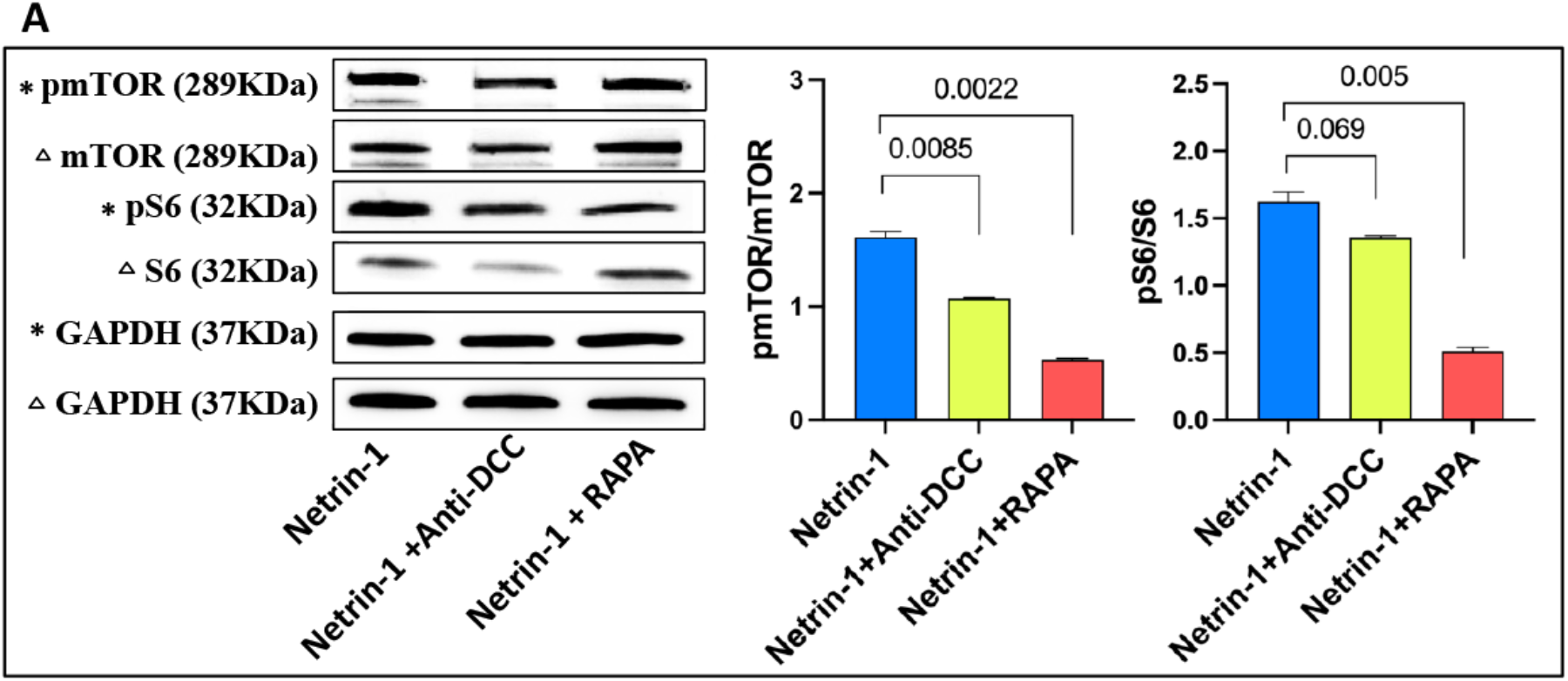
Netrin-1/DCC interactions recruit mTOR signaling: Western blot analysis was carried out on hRGCs^Brn3b-tDT^, preincubated with IgG/anti-DCC antibody, cultured in the presence of Netrin-1 and Netrin-1 plus rapamycin. The levels of pmTOR and pS6 decreased significantly when Netrin-1/DCC interactions were blocked or in the presence of rapamycin, compared to those in IgG controls suggesting the positive influence of Netrin-1/DCC interactions on mTOR signaling. Asterix and open triangles represent separate electrophoretic runs for the three treatment groups.

**Table S1:**
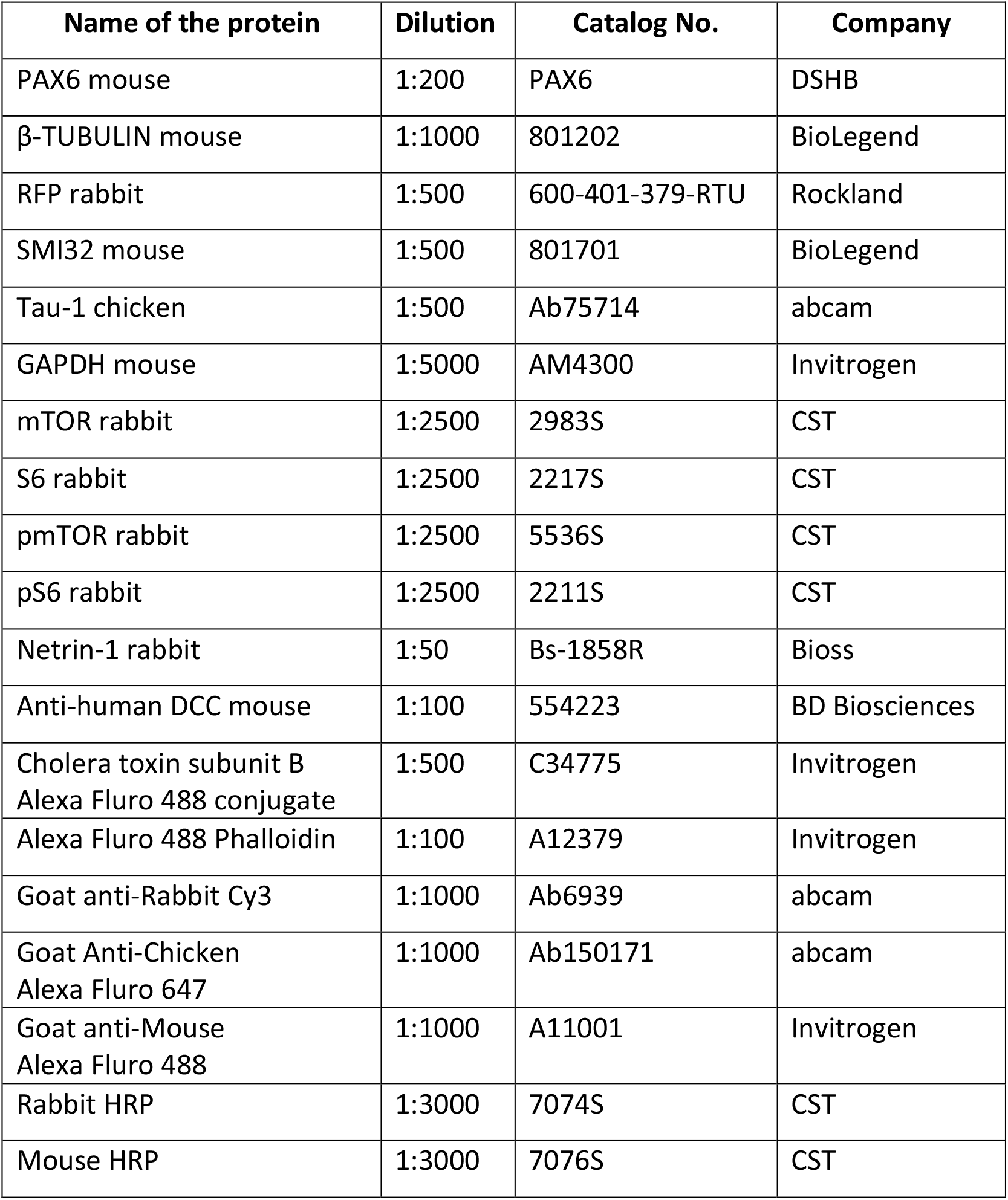
List of primary and Secondary Antibodies.

**Table S2:**
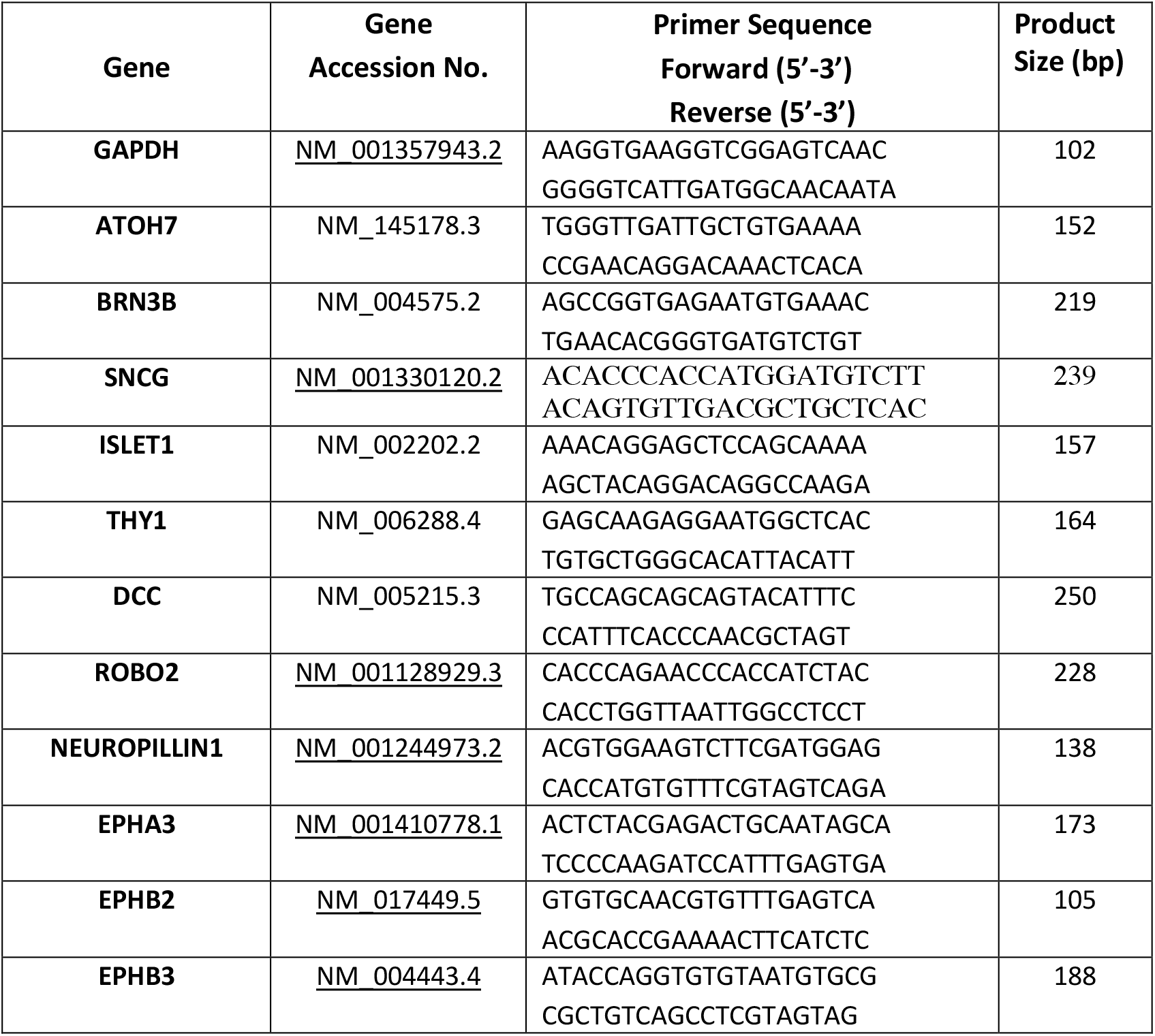
List of gene specific primers for Quantitative Real-Time PCR.

